# PathwaySeeker: Evidence-Grounded AI Reasoning over Organism-Specific Metabolic Networks

**DOI:** 10.64898/2026.04.14.718256

**Authors:** Lummy M. O. Monteiro, Niaz B. Chowdhury, Marjolein T. Oostrom, Jason E. McDermott, Kelly G. Stratton, Sutanay Choudhury, Jaydeep P. Bardhan

## Abstract

Metabolic activity is not an intrinsic property of an organism, but an emergent state shaped by environmental and experimental context. Despite recent advances in large language models (LLMs) and multi-omics profiling, current computational frameworks struggle to represent and reason over metabolism in a condition-specific manner. General-purpose AI systems operate on static, public biochemical knowledge, while multi-omics datasets capture dynamic measurements without a structured framework for mechanistic interpretation. As a result, metabolic networks remains analysis remains disconnected from the experimental states that define biological function. Here, we introduce PathwaySeeker, an evidence-grounded AI system for organism-specific metabolic network reasoning. PathwaySeeker reconstructs sample-specific metabolic graphs from integrated proteomic and metabolomic data, fine-tunes an LLM on the resulting graph structure, and verifies each reasoning step against the experimental graph through iterative hypothesis search, an approach we term Oracle-in-the-Loop inference. Every output claim carries explicit evidence provenance, distinguishing experimentally confirmed relationships from biochemically plausible hypotheses requiring validation. We demonstrate the system using multi-omics data from the non-model white-rot fungus *Trametes versicolor*, where PathwaySeeker recovers branched phenylpropanoid pathways and transparently stratifies confirmed reactions from testable extensions. Post-hoc thermodynamic analysis condition-specific metabolite dynamics support the biological feasibility of the reconstructed routes. By embedding experimental evidence provenance directly into language model-guided metabolic network reasoning, PathwaySeeker enables systematic differentiation between experimentally grounded knowledge and structured hypothesis, bridging frontier AI capabilities with organism-specific experimental evidence.

**Availability and Implementation:** PathwaySeeker is available at https://github.com/pnnl/PathwaySeeker

## Introduction

Metabolic pathways describe how living systems transform nutrient to biomass through coordinated enzyme-catalyzed reactions. While core metabolic processes are largely conserved across diverse environmental and physiological context, substantial portions of metabolism, including secondary pathways, branching points, and regulatory constraints are inherently conditional, with activity and connectivity shaped by environmental and experimental context.^1, 2^ Context-aware metabolic reasoning is therefore essential for elucidating biological function, identifying competing routes, and guiding metabolic engineering and bioprocess optimization.^3^ Despite substantial progress, current approaches exhibit three fundamental limitations. Curated pathway databases provide static, organism-agnostic representations that are largely decoupled from experimental context. Multi-omics datasets provide condition-specific molecular evidence but lack a structured reasoning layer to transform evidence into mechanistic pathway hypotheses. Meanwhile large language models, demonstrate advanced biochemical reasoning capabilities, ^4-6^ but do not inherently encode provenance, limiting their ability to discriminate experimentally supported relationships from plausible prior-driven inferences. We introduce PathwaySeeker, an AI system for condition-specific metabolic network reasoning that couples LLM-based inference with explicit verification against experimental knowledge. PathwaySeeker constructs context-specific knowledge graphs by mapping integrated proteomic and metabolomic measurements onto a structured metabolic reaction network, thereby annotating reactions and metabolites with condition-dependent evidence. The system is then fine-tuned on these evidence-grounded graphs and performs reasoning by constraining AI-generated hypotheses to graph-supported relationships. Each inferred pathway step is explicitly labeled as experimentally verified or as a testable hypothesis, providing transparent evidence provenance throughout the reasoning process.

Over decades, metabolic knowledge has been accumulated through experimental studies and organized into curated pathway databases.^7-10^ These resources provide canonical, organism-agnostic maps that serve an essential foundation for metabolic analysis. They also reflect historical biases toward well-studied organisms and conditions, and pathway annotations remain incomplete for many species, with condition-dependent routes and regulatory contexts underrepresented.^11, 12^ This limitation constrains discovery in non-model organisms ^2^ and restricts our ability to uncover novel biochemical capabilities beyond canonical pathways.

The growing availability of multi-omics data offers a path toward reconstructing metabolic activity directly from experimental evidence. Proteomics and metabolomics, in particular, provide complementary views of enzymatic potential and chemical transformations occurring within a given condition.^13^ Despite this promise, most computational approaches rely on individual omics layers, predefined organism-specific models, or association-based analyses that capture correlation without mechanism.^14-18^ As a result, they provide only partial insight into how metabolic systems reorganize across conditions ^2^ and cannot readily distinguish active pathways from latent enzymatic capacity (**Supplementary Table 1**).^19^

Recent efforts have integrated LLMs with multi-omics data. However, these approaches sidestep the central challenge of condition-specific metabolic inference: foundation models trained on multi-omics graphs emphasize signaling and regulatory networks rather than metabolism,^20^ metabolite-focused interfaces focus on chemical properties and annotations without integrating proteomic evidence into mechanistic reasoning,^21^ and agentic retrieval-augmented pipelines commonly query curated databases rather than learning to reason over experiment-derived metabolic graphs.^22^ As a result, none can distinguish pathways derived from experimental evidence from inferences drawn from the model’s training corpus.

We developed PathwaySeeker, an AI system that closes this gap by grounding language model reasoning in organism-specific experimental evidence. Importantly, this challenge cannot be resolved by applying standard adaptation strategies in isolation. Fine-tuning a language model on a multi-omics graph teaches it local graph structure but does not prevent the model from reverting to its pretraining priors when the graph is silent. Similarly, chain-of-thought prompting, 23 improves reasoning coherence but provides no mechanism for verifying intermediate claims against experimental data.

PathwaySeeker integrates graph-based fine-tuning and structured reasoning with a third element: iterative verification of each intermediate claim against a graph oracle. The oracle provides positive experimental evidence only, serving as a constraint rather than a generative source of knowledge. This verification loop adapts tree-structured and heuristic search paradigms that incorporate external feedback to guide LLM reasoning, ^24-26^ applying them to the setting of experimental metabolic networks. We term this approach Oracle-in-the-Loop inference. The oracle constrains reasoning to relationships supported by organism-specific proteomic and metabolomic measurements, while allowing hypothesis generation where experimental evidence is absent but connections are biologically plausible. By separating confirmed relationships from inferred extensions at runtime, every claim in the output remains transparently anchored to the experimental graph.

The system operates in three stages (**Fig. 1**). First, a modular pipeline constructs a condition-specific metabolic knowledge graph by projecting experimentally measured proteins and metabolites onto a curated reaction scaffold derived from KEGG. Reactions define the connections between compounds and enzymes, while proteomic and metabolomic measurements determine which nodes and edges are supported under the given experimental condition. Second, a schema-aware training data generator creates supervised examples from this graph, building on the Graph-aligned Language Model (GLaM) approach ^27^ with adaptations for metabolic semantics, teaching the model to distinguish experimentally verified relationships from inferred ones. Third, at inference time, an “Oracle-in-the-Loop search” orchestrates the fine-tuned model with a graph oracle that provides multi-omics relational evidence, applying step-level verification ^28^ against experimental data: the experimental graph confirms the presence of relationships but never treats absence as rejection, reflecting the inherent incompleteness of multi-omics-derived networks. A hypothesis-driven search iteratively generates candidate pathways maintaining only top-ranked candidates using a beam search ^29^ approach, checks each step against the graph oracle, and refines hypotheses based on retrieved evidence (**Fig. 2**).

**Fig 1.**
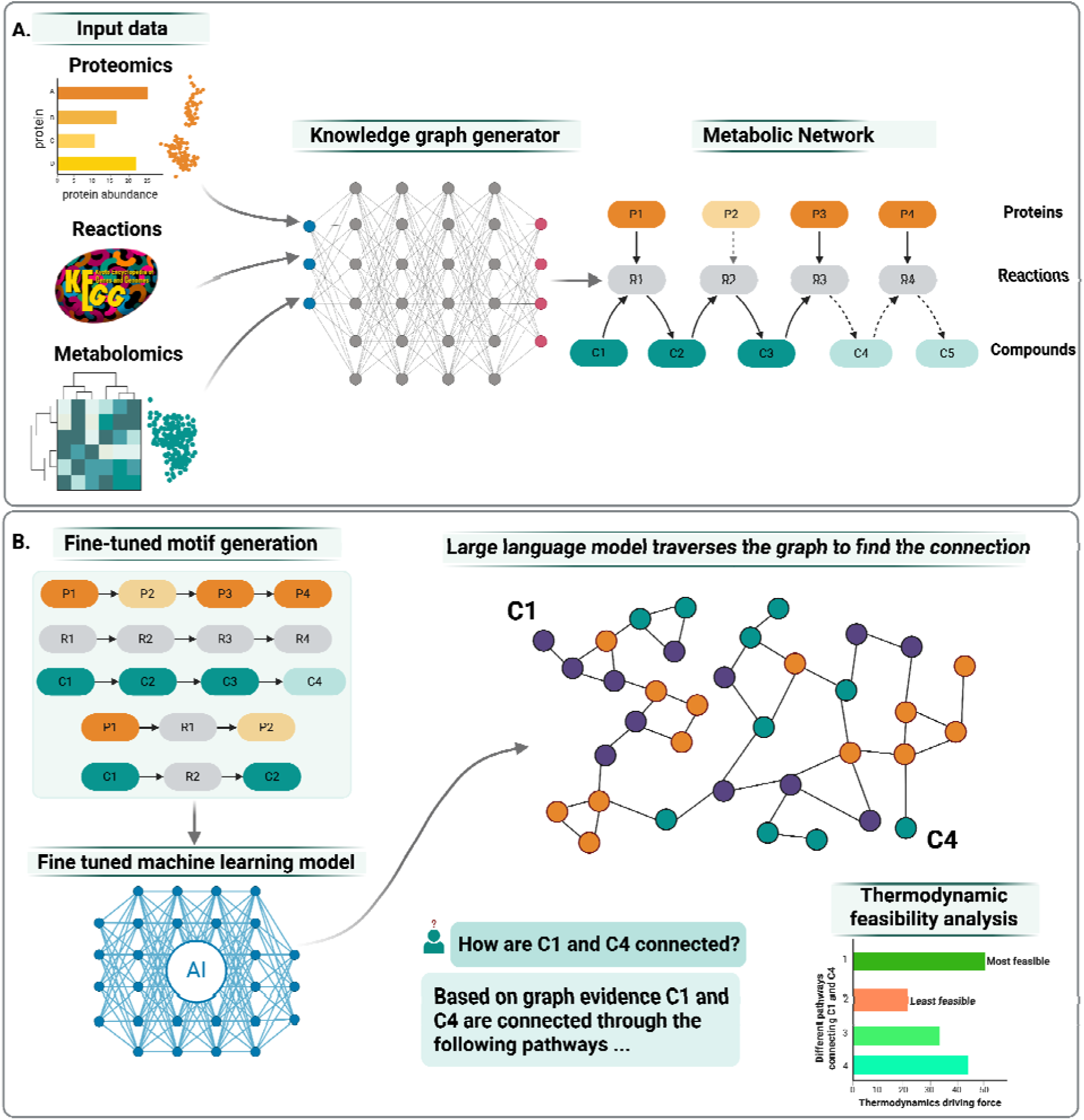
PathwaySeeker system overview. **A**. Condition-specific metabolic knowledge graph construction from integrated proteomic and metabolomic measurements. Detected proteins (P1-P4) are mapped to KEGG catalyzed reactions (R1-R4), which are linked to substrate and product compounds (C1-C5). Solid edges indicate relationships supported by experimental evidence; dashed edges indicate relationships inferred from a single omics layer. **B**. (Bottom left) Schema-aware training data generation extracts supervised examples at multiple scales: multi-step metabolic chains, shorter transformation sequences, and individual reaction units, each annotated with evidence type and confidence. (Bottom right) At inference time, the Oracle-in-the-Loop search iterates between the fine-tuned model, which generates candidate pathway hypotheses, and the experimental graph oracle, which verifies each reasoning step against observed evidence. Candidate pathways can be further filtered through thermodynamic feasibility analysis.

**Fig 2.**
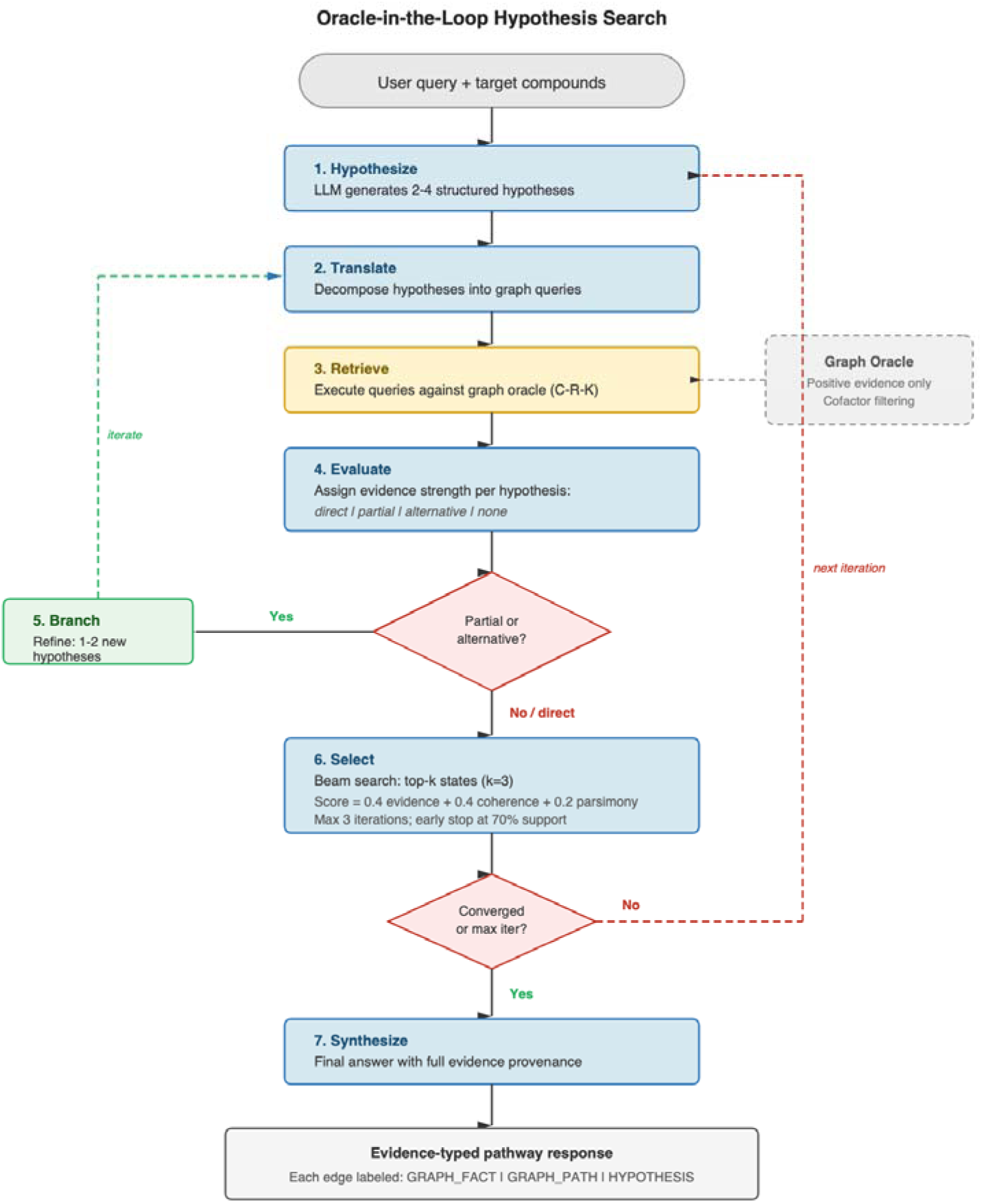
Oracle-in-the-Loop hypothesis search algorithm. Given a user query and target compounds, the fine-tuned model generates 2 to 4 structured hypotheses with compound, reaction, and enzyme identifiers (Stage 1). Hypotheses are decomposed into graph queries (Stage 2) and executed against a positive-evidence-only graph oracle that applies cofactor filtering to all path traversals (Stage 3). The model evaluates each hypothesis against retrieved evidence, assigning strength as direct, partial, alternative, or none, where none indicates absence from the experimental graph rather than biological implausibility (Stage 4). Hypotheses receiving partial or alternative support trigger generation of refined alternatives (Stage 5, green loop). A beam search selects the top-k candidate states (k = 3) by composite scoring of evidence support (0.4), biochemical coherence (0.4), and parsimony (0.2), iterating for a maximum of 3 rounds with early stopping at 70% or greater support (Stage 6, red loop). The final synthesis (Stage 7) produces a pathway response in which every edge carries explicit evidence provenance: GRAPH_FACT (verified single reaction), GRAPH_PATH (verified multi-step chain), or HYPOTHESIS (biochemically plausible, unverified, with confidence score and stated reasoning).

We demonstrate the reasoning system using multi-omics data from a non-model white-rot fungus *Trametes versicolor*, an ecologically and industrially relevant organism with well-documented lignin-degrading capabilities ^30-32^ but poorly characterized intracellular metabolism. In this setting, PathwaySeeker recovers known lignin-derived aromatic transformation routes and proposes condition-specific pathway extensions supported by experimental evidence. This case study demonstrates how evidence-grounded AI reasoning can uncover structured metabolic hypotheses directly from organism-specific datasets, extending beyond canonical pathway annotations.

## Results

The results section proceeds in three parts: construction and quantitative benchmarking of the condition-specific metabolic knowledge graph, evidence-typed pathway reasoning through phenylpropanoid case studies with biological and thermodynamic validation, and systematic evaluation across a large set of randomly sampled metabolic queries with comparison to ungrounded general-purpose models.

### Condition-specific Metabolic Knowledge Graph Construction for *Trametes versicolor*

To demonstrate the ability of PathwaySeeker to construct a condition-specific metabolic knowledge graph suitable for AI-guided reasoning, we focused on the white-rot fungus *Trametes versicolor*, an ecologically and industrially important non-model organism known for lignin and aromatic compound degradation ^30-32^ Despite its extensive use in lignin depolymerization studies, the intracellular metabolism of *T. versicolor* remains poorly characterized, owing to incomplete genome annotation, the absence of curated genome-scale metabolic models, and limited direct evidence of pathway activity. Consequently, intracellular metabolism in *T. versicolor* has often been interpreted through metabolic paradigm established in model organisms such as *Escherichia coli*, *Pseudomonas putida* or *Saccharomyces cerevisiae*,^11, 13, 31, 33^ rather than reconstructed from organism-specific experimental measurements. This indirect inference limits systematic discovery of condition-dependent pathways unique to *T. versicolor*.

In this study, we leveraged previously published multi-omics data for *T. versicolor* under defined experimental regimes, including variations in incubation mode (static versus agitation) and antioxidant supplementation, with cellobiose and 4-hydroxybenzoic acid as primary carbon sources.^13^ Rather than constructing separate graphs for each condition, we aggregated intracellular proteomic and metabolomic measurements across all cultivation settings to define the experimentally supported metabolic capacity of the organism. Proteins and metabolites detected in at least one condition were incorporated into the knowledge graph, thereby capturing the repertoire of biochemical functions observed under the tested experimental space. The dataset comprised intracellular proteomic and metabolomic measurements from biological replicates across these cultivation conditions. In total, 5859 proteins were quantified and 325 metabolites were detected across 4 experimental conditions, and these measurements were integrated for graph reconstruction.

The resulting knowledge graph captures metabolic activity as a directed reaction network in which nodes represent compounds and edges correspond to biochemical reactions, annotated by the type of experimental evidence supporting their inclusion (proteomics, metabolomics, or both; **Fig. 3**). We use an evidence-aware integration strategy that implements an inclusive, OR-based logic for network expansion rather than requiring simultaneous detection of enzymes and metabolites. From proteomics data, detected enzymes are mapped to their known catalytic reactions, and the corresponding substrates and products are incorporated into the network even when those compounds are not directly observed in the metabolomics dataset. Conversely, metabolomics data are used to identify reactions connecting detected compounds, independently of whether the catalyzing enzyme is observed at the protein level. As a result, reactions supported by metabolite-level evidence are retained in the graph even in the absence of detected enzymes, preventing gaps that would arise if edges were included only when both enzyme and metabolite evidence were present. This approach preserves network connectivity while maintaining explicit annotation of evidence type. This design enables recovery of plausible condition-specific metabolic routes in non-model organisms, where incomplete genome annotation and limited proteome coverage would otherwise obscure active pathways.

**Fig 3.**
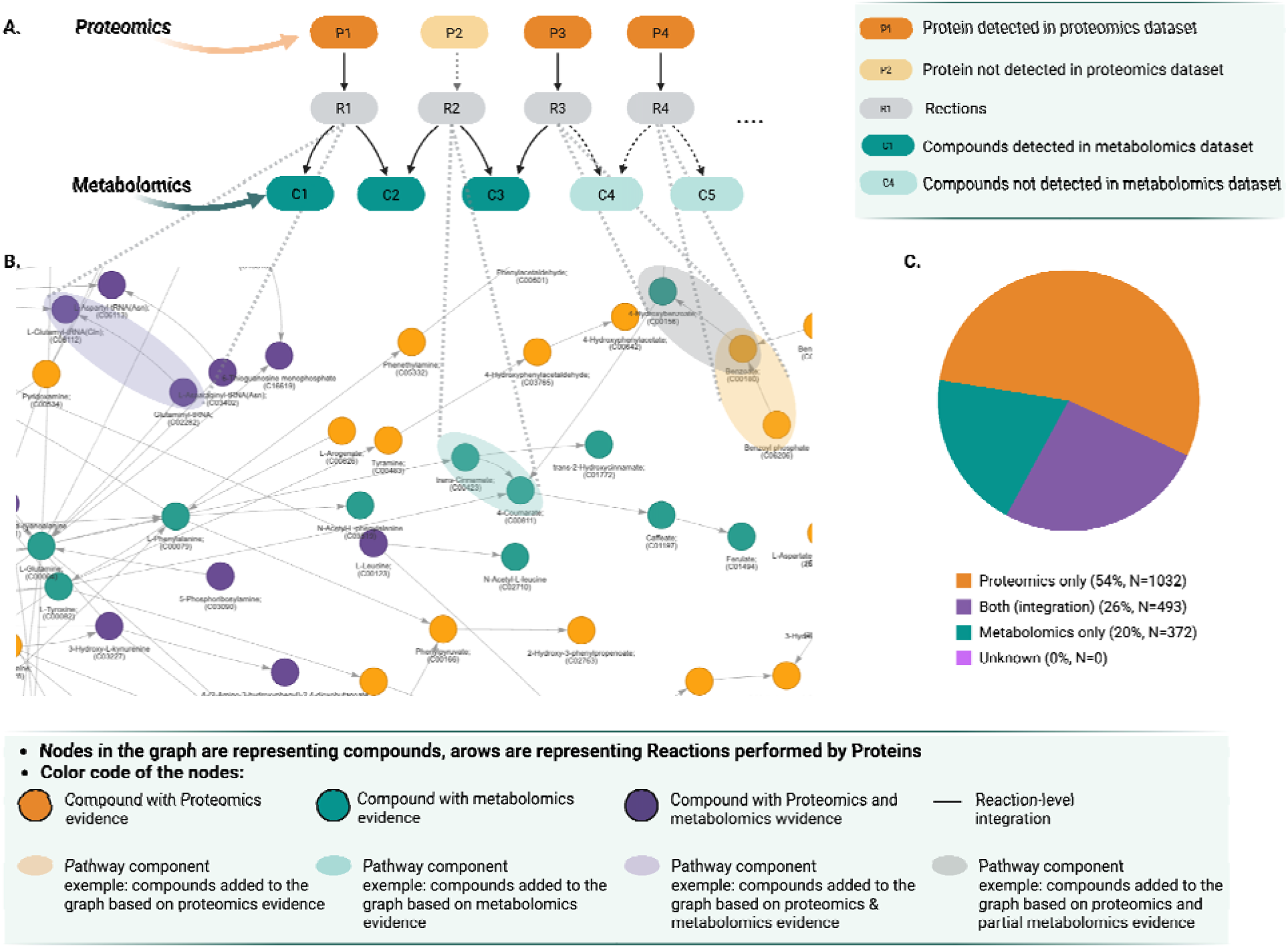
Condition-specific metabolic knowledge graph constructed from *T. versicolor* multi-omics data. **A.** Proteomics and metabolomics measurements are integrated into a directed three-layer graph of proteins, reactions, and compounds. Solid and dashed node borders distinguish entities detected in the corresponding omics dataset from those inferred through a single evidence layer. **B.** The reconstructed graph comprises 1,897 directed edges, colored by evidence source and displayed in the pie chart. **C.** proteomics only (54%), both omics layers (26%), or metabolomics only (20%).

### Evaluation of Knowledge Graph construction

Applying this knowledge graph construction approach to *T. versicolor* multi-omics datasets,^13^ we reconstructed a condition-specific metabolic knowledge graph comprising 3,402 unique reactions connecting 1,192 compounds through 1,897 directed edges. Reactions in the graph were cross-checked against KEGG annotations to confirm chemical and stoichiometric plausibility. This step functions as a sanity check to ensure that edges correspond to biochemically consistent transformations, rather than serving as an independent benchmark Integration of multi-omics evidence enabled annotation of each reaction based on the type of supporting data: 26% (493/1,897) were supported by both proteomics and metabolomics data, 54.4% by proteomics alone, and 19.6% by metabolomics alone (**Supplementary Table 2**). 163 of 181 experimentally detected metabolites were successfully mapped to the graph, corresponding to 90.1% compound coverage (**Supplementary Table 3**).

To assess the reliability and biological fidelity of the reconstructed graph, we first performed a sanity check against KEGG references to ensure that all reactions inferred in *T. versicolor* correspond to biochemically plausible transformations and that enzyme–substrate mappings were correctly established. This step does not constitute an independent validation but confirms that the graph does not contain chemically impossible reactions or mapping errors. We then assessed the recovery of known reactions using curated references in two complementary ways: comparison to the global KEGG reaction universe (12,312 reactions) evaluates precision and guards against false-positive inference, while comparison to the well-annotated model organism *Saccharomyces cerevisiae* (1,512 reactions) assesses the recovery of eukaryotic metabolic capabilities. Relative to the global KEGG universe, PathwaySeeker achieved a recall of 27.6% reflects intentional restriction to reactions supported by experimental evidence in *T. versicolor*. Relative to *S. cerevisiae*, recall increased to 72.6%, with a precision of 0.323, reflecting the tradeoff between recovering broadly conserved eukaryotic metabolic functions and maintaining organism and condition specific inference (**Supplementary Table 4**). Together, these results indicate that PathwaySeeker’s knowledge graph construction recovers biologically relevant reactions while maintaining conservative, organism-consistent inference.

### Enabling Evidence-Guided Reasoning over a Multi-Omics Knowledge Graph

The reconstructed metabolic graph provides a condition-specific biochemical scaffold, but its true utility lies in enabling structured reasoning over experimental evidence. This is enabled by the fact that every claim in PathwaySeeker’s output carries explicit evidence provenance. Relationships verified in the experimental graph are labeled GRAPH_FACT (individual reactions) or GRAPH_PATH (multi-step pathways where all edges are experimentally confirmed). Connections that are biochemically plausible but extend beyond the current graph annotations are labeled HYPOTHESIS, with stated biochemical reasoning and confidence scores.

This evidence typing serves a deeper purpose than provenance annotation. PathwaySeeker is not restricted to reasoning within the experimental graph. It draws on the full biochemical knowledge embedded in the underlying language model, but maintains a clear boundary between what the experiment confirms and what remains a hypothesis. The key design principle is asymmetric treatment of graph evidence: presence in the multi-omics graph provides positive confirmation, but absence does not constitute rejection. The *T. versicolor* graph comprises 19 disconnected subgraph components across approximately 4,000 nodes, with fragmentation arising from gaps in experimental annotation rather than true metabolic isolation. Rather than concluding "no path exists" biologically when the graph has no evidence for a connection, PathwaySeeker generates testable hypotheses while clearly marking them as unverified, enabling researchers to identify exactly which pathway steps require experimental follow-up.

### Recovering experimentally grounded pathways beyond the AI training cutoff

We fine-tuned PathwaySeeker on OpenAI’s GPT-4.1as the baseline since it was the only modeled allowed by Azure OpenAI’s Fine-Tuning API, whose training cutoff (June 2024) predates the publication of Monteiro et al. (2025).^13^ We then posed phenylpropanoid pathway queries that Monteiro et al. had investigated through manual curation of the same *T. versicolor* multi-omics dataset. Any recovery that matches the manually curated result must therefore derive from experimental grounding rather than memorization of the published literature, providing a direct test of whether PathwaySeeker can independently reconstruct branched, multi-step pathways with transparent evidence provenance.

Phenylpropanoid metabolism underlies lignin-derived aromatic transformation in white-rot fungi and defines how depolymerized carbon enters central metabolism. In *T. versicolor*, Monteiro et al. (2025) manually reconstructed this pathway by integrating proteomic and metabolomic evidence with homology-based inference from reference organisms including *Arabidopsis thaliana*, *Pseudomonas putida*, and *Saccharomyces cerevisiae*.^13^ Their analysis identified a canonical backbone from L-phenylalanine to ferulate, convergence with an alternative L-tyrosine entry route, and branching toward 4-hydroxybenzoate. Because a curated metabolic model was unavailable, several intermediate steps were incorporated through cross-species homology to construct a mechanistically coherent hypothesis. We therefore asked whether PathwaySeeker could independently recover this branched phenylpropanoid architecture directly from the experimentally constructed knowledge graph, without homology-driven gap filling, pathway templates, or access to post-2024 literature. Specifically, we evaluated its ability to reconstruct (i) the canonical backbone, (ii) entry-point convergence, and (iii) branch formation within the pathway topology.

When queried about the metabolic connection between L-phenylalanine (C00079) and ferulate (C01494), PathwaySeeker recovered the complete canonical phenylpropanoid backbone: L-phenylalanine → trans-cinnamate (via R00697) → 4-coumarate (via R02253) → caffeate (via R07826) → ferulate (via R03366), with all four reactions and five intermediates verified as GRAPH_PATH in the experimental graph (**Fig. 4A, left**). Beyond this verified core, PathwaySeeker identified alternative entry points into the pathway, noting that 4-coumarate can also be produced from L-tyrosine and 4-hydroxybenzoate.

**Fig 4.**
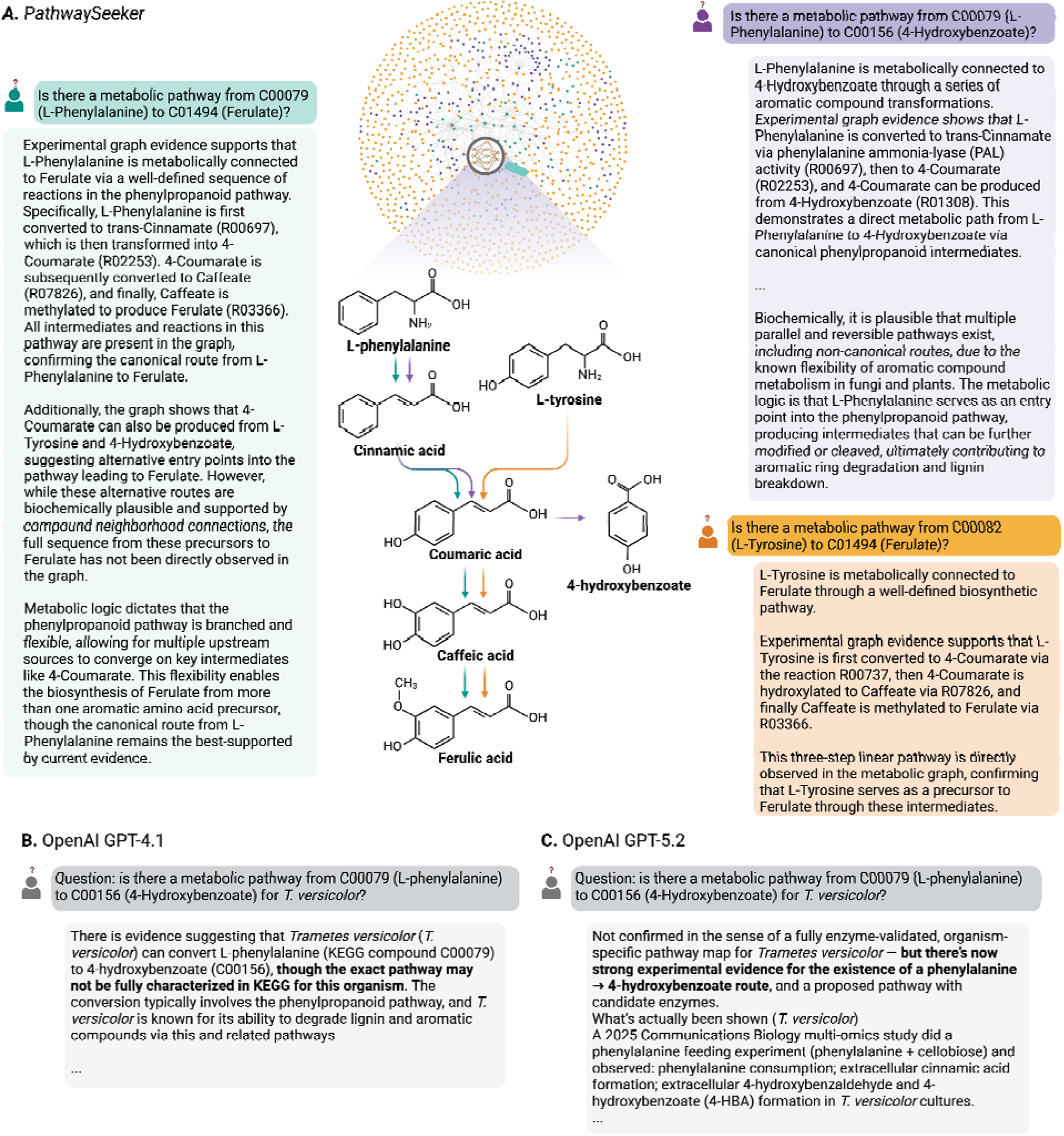
Evidence-typed reasoning versus general-purpose language models. GPT-4.1’s training cutoff (June 2024) predates Monteiro et al. (2025),^13^ so PathwaySeeker’s recovery of *T. versicolor* phenylpropanoid pathways derives from experimental grounding, not memorization. (**A**) PathwaySeeker responses illustrating two discovery modes. Confirmation: the canonical phenylalanine-to-ferulate backbone (left-green) and tyrosine-derived alternative entry (bottom right-orange) are verified as active in *T. versicolor* under the measured conditions. Divergence: the branch from coumarate to 4-hydroxybenzoate (top right-purple) emerges directly from the experimental graph, recovering a route that model-organism-guided manual curation did not prioritize. (**B**) GPT-4.1 produces biochemically accurate but organism-agnostic responses with no evidence provenance. (**C**) GPT-5.2 returns correct pathways but cannot distinguish the user’s experimental evidence from pretraining knowledge.

A second query revealed PathwaySeeker’s treatment of pathway branching. The model identified the phenylpropanoid backbone from phenylalanine through trans-cinnamate and 4-coumarate, with 4-coumarate serving as a branch point toward 4-hydroxybenzoate (C00156) via R01308 (**Fig. 4C, top right**). PathwaySeeker distinguished this graph-verified branch from additional biochemically plausible parallel routes, noting the known flexibility of aromatic compound metabolism in fungi and plants while clearly marking these extensions as hypotheses. This result is particularly notable because the route from p-coumarate to 4-hydroxybenzoate emerged directly from the experimental graph structure, without invoking intermediates from cross-species homology transfer, in contrast to the reconstruction approach taken by Monteiro et al. (2025) based on enzymatic machinery from *Pseudomonas putida* and *Rhodosporidium toruloides*.^13^

A third query confirmed the alternative entry point. When asked about the connection between L-tyrosine (C00082) and ferulate (C01494), PathwaySeeker recovered a verified three-step pathway: L-tyrosine → 4-coumarate (via R00737) → caffeate (via R07826) → ferulate (via R03366), converging with the phenylalanine-derived route at 4-coumarate and continuing through shared downstream reactions (**Fig. 4A, bottom right**). The independent recovery of this convergent architecture, without manual curation or pathway templates, validates PathwaySeeker’s ability to discover branched metabolic topology directly from experimental evidence.

In each of these three cases, PathwaySeeker’s responses matched the manually curated pathway architecture while providing two capabilities unavailable through manual curation: automatic evidence provenance for every pathway step, and explicit generation of testable hypotheses for pathway extensions beyond the current experimental graph.

The preceding results demonstrate PathwaySeeker’s core reasoning loop. The following two analyses apply independent, post-hoc validation to the system’s outputs. We first assess whether predicted pathways are thermodynamically feasible. We then evaluate whether their constituent enzymes and metabolites exhibit coordinated, condition-dependent abundance changes consistent with active pathway engagement across experimental treatments. Neither analysis is currently integrated into the Oracle-in-the-Loop search; both serve as external checks on the quality of its predictions and could be incorporated as scoring functions within the beam search in future iterations.

### Thermodynamic validation of pathway directionality and feasibility

Before evaluating biological activity, predicted pathways were assessed for thermodynamic feasibility. Because network-based and AI-driven inference can generate multiple candidate biochemical routes, PathwaySeeker outputs were further filtered through thermodynamic feasibility analysis using eQuilibrator 3.0.^34^ Candidate pathways were evaluated using Max-min Driving Force (MDF) analysis to calculate the overall pathway driving force. Change in Gibbs free energy of each reaction was calculated with standard eQuilibrator parameters: pH 7.0, ionic strength 250 mM, pMg 3.0, and metabolite concentrations constrained between 0.00001 and 10 mM. Pathways with positive overall MDF values were considered thermodynamically feasible and retained for further analysis, whereas routes exhibiting negative driving force were flagged as infeasible under the tested constraints. Thermodynamic filtering thereby distinguished energetically robust pathways from alternatives incompatible with intracellular constraints. This independent validation confirms which of PathwaySeeker’s graph-grounded inferences are thermodynamically plausible and demonstrates that evidence-typed AI reasoning can be coupled with physics-based filtering to prioritize experimentally testable hypotheses (**Fig. 5**).

**Figure 5.**
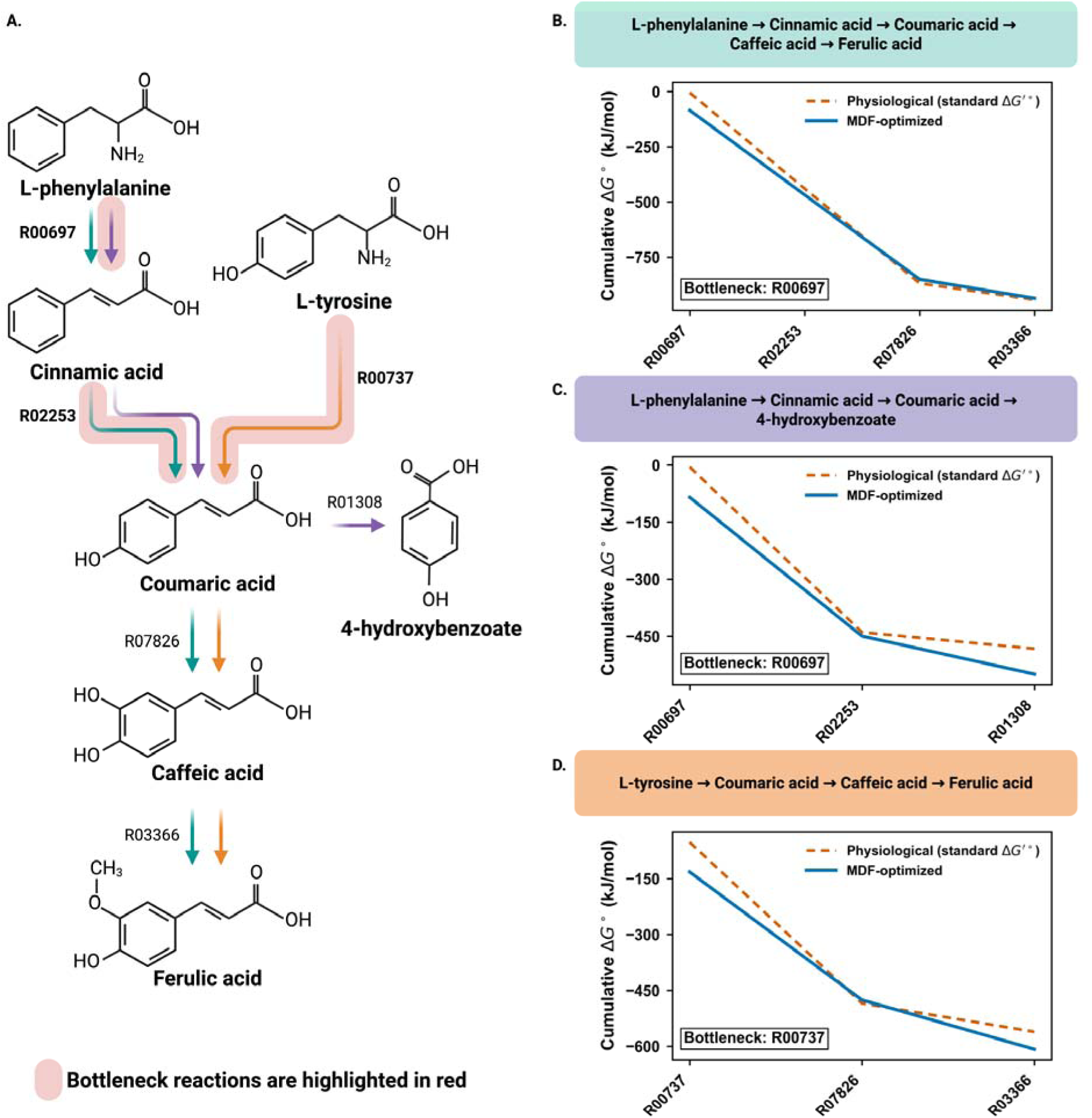
Thermodynamic validation and bottleneck identification in AI-inferred aromatic pathways. **A.** Graph-grounded reconstruction of three AI-inferred pathways in T. versicolor: phenylalanine-to-ferulate, tyrosine-to-ferulate, and phenylalanine-to-4-hydroxybenzoate. Reactions identified as thermodynamic bottlenecks are highlighted in red. **B–D.** Max–min Driving Force (MDF) analysis of each pathway using eQuilibrator 3.0 under standard biochemical conditions (pH 7.0, ionic strength 250 mM, pMg 3.0; metabolite concentration constrained between 10 and 10 mM). Cumulative Gibbs free energy (ΔG′) profiles are shown for both physiological standard ΔG′° and MDF-optimized conditions. All three pathways exhibit an overall favorable free energy landscape, while specific reactions impose local thermodynamic constraints (bottlenecks), consistent with those highlighted in panel A. Together, these results demonstrate that PathwaySeeker’s graph-grounded AI inference generates thermodynamically plausible routes and enables systematic identification of rate-limiting steps for experimental prioritization.

Among the AI-inferred candidate pathways in *T. versicolor*, thermodynamic analysis distinguished energetically favorable routes from alternatives containing unfavorable steps (**Fig. 5**). Step-level ΔG′ values derived from the MDF optimization further identified reactions operating with minimal driving force within otherwise feasible pathways. These minimal-driving-force steps represent thermodynamic bottlenecks, as they constrain the overall pathway flux under the tested intracellular conditions. This layered approach, structural inference followed by thermodynamic filtering, provides a principled mechanism to prioritize pathway hypotheses and identify rate-limiting steps that may represent targets for metabolic engineering.

### Biological validation through condition-specific metabolite abundance profiling

Following thermodynamic validation, we evaluated whether the inferred pathways exhibited signatures of functional engagement across experimental conditions. Importantly, while multi-omics data were used to construct the structural knowledge graph (i.e., to establish which enzymes and metabolites were present and potentially connected), no information about condition-dependent abundance changes was incorporated into the pathway search itself. Here, we introduce an orthogonal evaluation layer by overlaying quantitative metabolite abundance profiles onto the predicted pathway subgraphs to assess coordinated, condition-specific dynamics. This analysis therefore evaluates dynamic coherence rather than structural plausibility, distinguishing metabolic potential (graph connectivity) from metabolic activity (coordinated abundance variation).

Applied to the phenylpropanoid backbone in *T. versicolor*, this integration revealed coordinated abundance shifts across cultivation conditions in key intermediates, including trans-cinnamate, 4-coumarate, and caffeate (**Fig. 6**). The convergence nodes such as 4-coumarate displayed condition-dependent variation consistent with their role as metabolic branch points verified in the experimental graph. In contrast, metabolites associated with the 4-hydroxybenzoate branch exhibited abundance profiles distinct from upstream intermediates, suggesting differential partitioning of aromatic flux across environmental contexts. This abundance-aware representation enables discrimination between structural connectivity and condition-specific metabolic activity.

**Figure 6.**
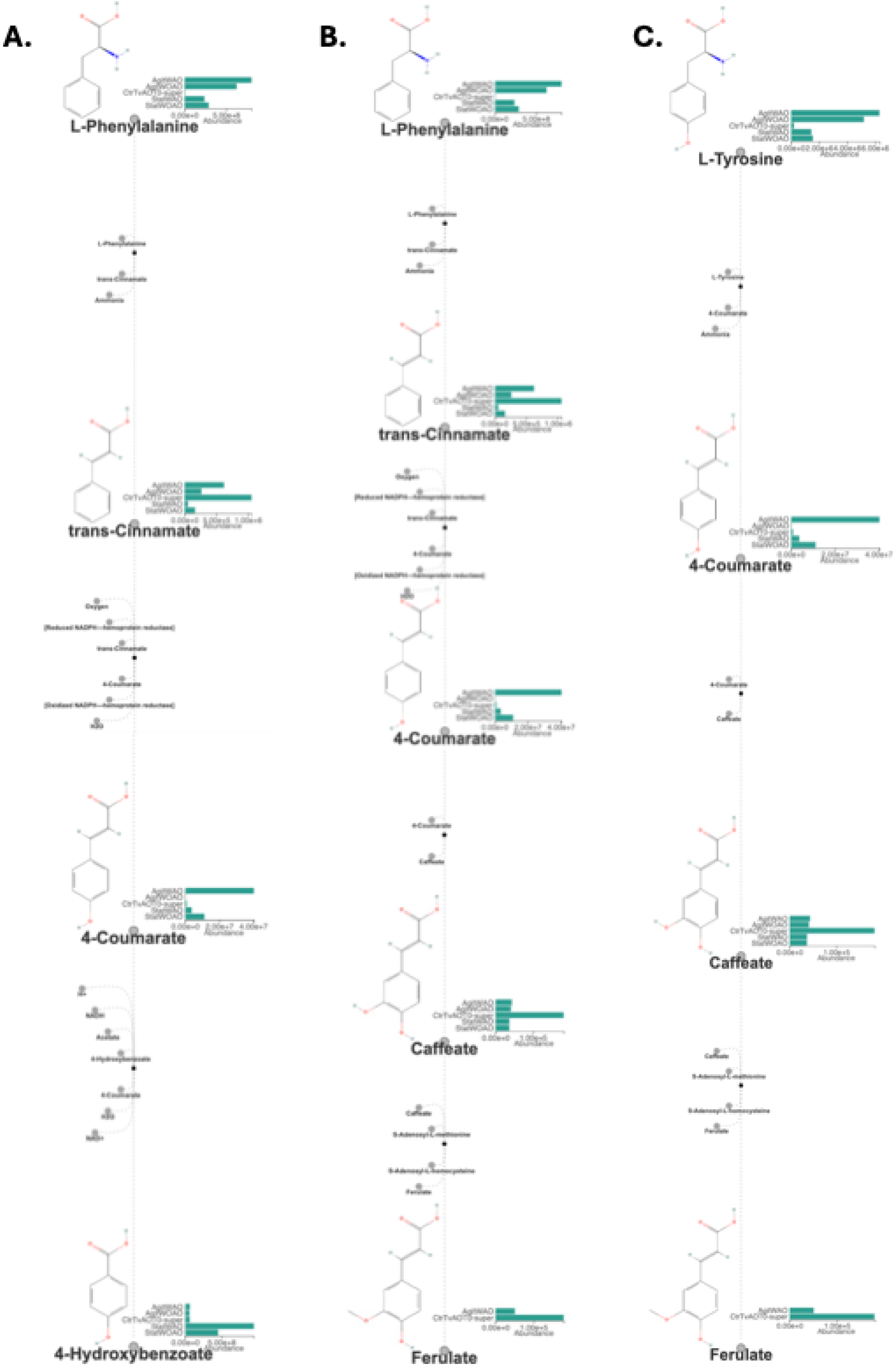
Condition-resolved metabolite dynamics validate active pathway utilization. **A–C**. Metabolite abundances overlaid onto three PathwaySeeker-inferred aromatic pathways in *T. versicolor*: phenylalanine-to-4-hydroxybenzoate, phenylalanine-to-ferulate, and tyrosine-to-ferulate. Each pathway is shown with full reaction context, including cofactors, alongside condition-specific abundance profiles for all intermediates. Pathway segments annotated as GRAPH_PATH display coordinated metabolite shifts across cultivation conditions, consistent with active metabolic utilization. In contrast, HYPOTHESIS branches lack coherent abundance dynamics, indicating condition-dependent inactivity or unrealized flux. This integration distinguishes structural connectivity from condition-specific metabolic activity, linking graph topology to experimentally observed compound-level dynamics.

Pathway segments verified as GRAPH_PATH show coherent metabolite dynamics across conditions; HYPOTHESIS extensions that lack such support become concrete targets for experimental follow-up. This closes the loop from computational prediction to experimental prioritization, which is the practical value of evidence-typed reasoning for non-model organisms where experimental resources are limited. Together, these results demonstrate that PathwaySeeker not only reconstructs structurally and thermodynamically plausible pathways, but also identifies condition-resolved metabolic activity patterns that refine experimental prioritization in non-model systems.

### Systematic evaluation of evidence provenance and scientific quality

To quantify PathwaySeeker’s performance beyond targeted case studies, we evaluated the system on 64 metabolic queries spanning connected and disconnected regions of the *T. versicolor* graph, including the four phenylpropanoid queries described above (**Table 1**).

**Table 1.** Systematic evaluation of PathwaySeeker across 64 metabolic queries in *T. versicolor*. Queries span connected compound pairs (both endpoints in the same graph component), unconnected pairs (endpoints in different components), and phenylpropanoid case studies manually curated by domain experts. Experimental Evidence Ratio (EER) measures the fraction of reported pathway edges verified in the organism-specific multi-omics graph. Scientific quality was assessed by an independent LLM judge (GPT-4.1) on four dimensions (1 to 5 scale). Five of 64 queries produced errors and are excluded.

| Query Category | N | EER (%) | Scientific Reasoning | Specificity | Evidence Transparency | Clarity | Overall Quality |
| --- | --- | --- | --- | --- | --- | --- | --- |
| Connected | 37/40 | 40.8 | 5.00 | 4.76 | 4.97 | 5.00 | 4.95 |
| Unconnected | 18/20 | 0.9 | 4.78 | 3.44 | 5.00 | 4.94 | 4.44 |
| Phenylpropanoid | 4/4 | 8.4 | 5.00 | 4.50 | 5.00 | 4.75 | 4.75 |
| All queries | 59/64 | 26.4 | 4.93 | 4.34 | 4.98 | 4.97 | 4.78 |

We introduce the Experimental Evidence Ratio (EER) as a provenance characterization metric: the fraction of pathway edges in each response that are directly confirmed by the organism-specific multi-omics graph. EER measures the composition of each response, the balance between experimentally grounded claims and novel hypotheses generated through biochemical reasoning, rather than serving as an accuracy score. A lower EER does not indicate lower quality; it indicates that the model is reasoning beyond the boundaries of the experimental graph, which is the intended behavior when graph annotations are incomplete.

PathwaySeeker achieved an EER of 26.4% across 64 queries, indicating that approximately one quarter of reported pathway edges are experimentally verified in the *T. versicolor* graph while three quarters represent explicitly labeled hypotheses extending beyond the graph. Scientific quality was assessed independently through an LLM-based judge evaluating four dimensions: scientific reasoning (biochemical logic and mechanistic depth), specificity (use of concrete KEGG identifiers), evidence transparency (distinction between verified and hypothesized claims), and clarity. PathwaySeeker achieved an overall quality score of 4.78 out of 5, with the highest scores on evidence transparency and scientific reasoning (**Table 1**), confirming that the system provides biochemically sound, well-structured responses with clear evidence provenance.

## Discussion

General-purpose language models encode biochemical knowledge, yet this knowledge is inherently organism-agnostic. They assign plausibility to metabolic claims conditioning on whether specific reactions are active under defined context. In contrast, multi-omics measurements provide direct molecular evidence of enzymes and metabolites present in a biological system, but lack an inferential framework for interpreting that evidence in the context of broader biochemical knowledge. PathwaySeeker couples these two information sources through evidence typing, producing outputs in which every claim is annotated with its epistemic status relative to the experimental graph. This coupling gives rise to four distinct modes of knowledge acquisition, each illustrated by our results in *T. versicolor*.

PathwaySeeker operates across distinct epistemic regimes that together reshape how metabolic knowledge is evaluated. In its most direct form, general biochemical plausibility becomes experimentally grounded activity. The recovery of the complete phenylpropanoid backbone from L-phenylalanine to ferulate as GRAPH_PATH, with all reactions and intermediates supported by proteomic and metabolomic evidence, illustrates how canonical metabolism is translated from expectation into organism- and condition-specific confirmation. Although this pathway is well established in plant and fungal systems, PathwaySeeker provides an organism- and condition-specific verification that it is active in *T. versicolor* under the measured cultivation conditions. The temporal gap between GPT-4.1’s training cutoff (June 2024) and the publication of Monteiro et al. (2025)^13^ ensures that this verification derives from experimental grounding rather than memorization of the published result.

Beyond confirming canonical pathway presence, PathwaySeeker also enables functional stratification within structurally supported networks. In *T. versicolor*, structural confirmation of the phenylpropanoid backbone did not uniformly translate into functional engagement across cultivation conditions. While the canonical L-phenylalanine → ferulate route was fully supported as GRAPH_PATH, quantitative metabolite profiling revealed differential abundance patterns along its branches. Intermediates such as trans-cinnamate, 4-coumarate, and caffeate exhibited coordinated condition-dependent shifts, consistent with active flux through the backbone. In contrast, metabolites associated with the 4-hydroxybenzoate branch displayed distinct abundance dynamics, suggesting context-dependent attenuation or rerouting of aromatic carbon under specific cultivation regimes. Thermodynamic profiling further refined this picture. Although the overall backbone satisfied positive MDF constraints, step-level analysis identified reactions operating with minimal driving force, highlighting specific energetic bottlenecks within otherwise feasible routes. Together, these findings demonstrate that topological connectivity alone is insufficient to infer metabolic activity: abundance coherence distinguishes dynamically engaged segments, while MDF constraints identify energetically limiting steps. This layered interpretation transforms pathway reconstruction from structural mapping into condition-resolved functional stratification.

In addition to stratifying activity within confirmed pathways, experimentally grounded reasoning can also reshape pathway topology itself. More unexpectedly, experimentally constrained inference can diverge from model-organism-guided reconstructions. The route from p-coumarate to 4-hydroxybenzoate via R01308 emerged directly from the experimental graph topology, without reliance on cross-species homology transfer as in the manual reconstruction by Monteiro et al. (2025).^13^ In contrast, manual reconstruction required importing reactions from well-characterized reference organisms to bridge gaps in the *T. versicolor* dataset. By restricting inference to experimentally observed molecules and enzymes, PathwaySeeker identified an alternative reaction organization that would not typically be prioritized when reasoning is filtered through canonical architectures. This divergence highlights how model-organism bias can shape pathway reconstruction and demonstrates that graph-grounded reasoning may expose metabolically coherent routes that remain underexplored when inference is anchored to familiar reference species. (**Fig 4 – top right-purple**).

Together, confirmation and divergence define only part of the system’s epistemic landscape; the third regime emerges where evidence is incomplete. Finally, a substantial portion of the reconstructed network consists of biochemically plausible but yet unconfirmed connections. HYPOTHESIS edges, which constitute approximately three-quarters of PathwaySeeker’s reported pathway edges (EER = 26.4% across 64 queries), represent biochemically plausible connections carrying stated reasoning and confidence scores that are not yet confirmed by the experimental graph. These are not incidental byproducts of the reasoning process; they are the system’s principal output for driving experimental follow-up. We demonstrate the infrastructure for this mode but leave experimental confirmation as future work, as biochemical validation of individual pathway predictions requires targeted enzyme assays, isotope tracing, and gene deletion experiments operating on a long timescale.

At a systems level, these epistemic distinctions are reflected in a quantifiable contraction of biochemical uncertainty. Experimental grounding imposes a measurable reduction in biochemical uncertainty. This reduction is quantifiable. GPT-4.1’s training corpus encompasses all 12,312 KEGG reactions; the *T. versicolor* experimental graph retains 3,402 (27.6%). Of the approximately 1.3 million possible non-cofactor compound pairs derivable from the graph’s 1,153 backbone metabolites, only 2,831 (0.21%) are connected by verified metabolic paths. A general-purpose LLM treats the full combinatorial space as potentially valid; the experimental graph eliminates 99.79% of those candidate connections from consideration. The principal informational contribution of experimental grounding is therefore not the introduction of novel reactions, but the systematic contraction of biochemical plausibility into an observed, organism-specific subspace. In information-theoretic terms, experimental data acts as a powerful prior that collapses the combinatorial search space.

This narrowing operates at the level of individual pathway queries. For example, the connection between L-phenylalanine (C00079) and ferulate (C01494) presents multiple biochemically defensible routes in an unconstrained metabolic universe. Within the experimentally grounded graph, however, this multiplicity is reorganized: one route is confirmed, partial evidence supports a limited subset of alternatives, and the remainder are explicitly labeled as hypotheses with graded confidence. The output is not merely filtered biochemical knowledge, but a structured representation in which every claim is accompanied by provenance and evidential status, enabling differentiated experimental action.

Maintaining this structured uncertainty requires architectural safeguards that prevent epistemic collapse. PathwaySeeker explicitly distinguishes between hypotheses and hallucinations, ensuring transparent representation of uncertainty. Hypotheses are unverified claims assigned confidence scores, supported by biochemical reasoning, and consistent with known metabolic constraints. Hallucinations, in contrast, are claims presented as verified that contradict evidence or violate constraints. Architectural safeguards enforce this distinction: evidence typing prevents unverified claims from being misinterpreted as confirmed; cofactor-constrained heuristics eliminate biologically implausible hub-mediated shortcuts; and the positive-evidence-only oracle ensures that absence in the graph is never treated as biological impossibility. While incorrect hypotheses may still be generated, they remain auditable and traceable against the experimental graph, providing an operationally meaningful standard for uncertainty.

This design places PathwaySeeker within a distinct methodological position relative to existing AI–omics integration frameworks. Compared to existing approaches, PathwaySeeker occupies a distinct methodological space. mosGraphGPT^20^ trains foundation models on multi-omics signaling graphs but does not separate experimental evidence from pretraining priors and focuses on regulatory rather than metabolic networks. MetaboliteChat^21^ offers multimodal analysis of metabolites but operates on chemical properties without integrating proteomic evidence into mechanistic reasoning. Agentic RAG pipelines, such as Arowolo et al.,^22^ query curated pathway databases at inference but cannot learn from experiment-derived graph structures or differentiate database-sourced from model-inferred claims. In contrast, PathwaySeeker trains directly on experimentally reconstructed metabolic graphs, performs heuristic search with runtime verification, and annotates every edge with its evidence type and provenance. Its Oracle-in-the-Loop inference pattern, in which AI-generated hypotheses are iteratively checked against a positive-evidence-only oracle, is broadly applicable to any domain combining an incomplete but reliable evidence graph with a reasoning-capable language model.

While the primary analyses focus on a single organism under defined cultivation conditions, the underlying architecture is organism-agnostic. The schema-aware training pipeline, evidence typing protocol, and cofactor constraints were successfully applied to reconstruct metabolic knowledge graphs from independent multi-omics datasets in additional species (Results in the GitHub repository).^35^ However, a systematic cross-organism evaluation of reasoning performance has not yet been conducted. The extent to which graph density, fragmentation patterns, and metabolic domain influence confirmation rates and hypothesis quality remains to be quantitatively characterized across organisms. Addressing these variables through comparative multi-species deployment will clarify how biological coverage shapes system performance.

Second, none of the HYPOTHESIS edges have been experimentally validated. While phenylpropanoid case studies demonstrate recovery of canonical pathways and generation of plausible extensions, the scientific value of these extensions ultimately depends on targeted experimental testing. High-priority candidates, such as edges connecting aromatic intermediates to central carbon metabolism in *T. versicolor*, should be validated via enzyme assays or isotope tracing.

Third, the evaluation framework currently combines automated oracle metrics, LLM-as-judge assessment, and blinded expert review, but lacks systematic failure-mode analysis. Understanding the conditions under which PathwaySeeker generates low-quality hypotheses whether due to graph sparsity, ambiguous metabolic semantics, or base model limitations is essential for establishing principled confidence thresholds and guiding experimental investment. Future directions include multi-organism deployment, integration with transcriptomic and flux data, and systematic characterization of failure modes.

In summary, PathwaySeeker demonstrates that coupling large language model reasoning with experimentally reconstructed knowledge graphs transforms metabolic inference from plausibility estimation into evidence-stratified discovery. Rather than replacing biochemical expertise, the framework reorganizes it: canonical pathways become organism-specific confirmations, structurally intact routes are dynamically and thermodynamically stratified, and unverified connections are converted into auditable, prioritized hypotheses. By contracting the combinatorial metabolic search space into an experimentally grounded subspace, PathwaySeeker operationalizes uncertainty in a way that is transparent, quantifiable, and experimentally actionable. This architecture is not limited to metabolism, but exemplifies a broader paradigm in which AI systems reason within, rather than beyond, the constraints of empirical evidence.

## Methods

The modular architecture of PathwaySeeker, integrating graph-grounded reconstruction with AI-guided hypothesis expansion, is summarized in **Supplementary Fig. 1**. PathwaySeeker operates in three stages (**Fig. 1**): condition-specific metabolic graph construction from integrated multi-omics data (**Fig. 1A**), schema-aware training data generation and model fine-tuning (**Fig. 1B**), and Oracle-in-the-Loop inference with hypothesis-driven beam search (**Fig. 1C**).

### Detailed pipeline implementation

PathwaySeeker is implemented as a modular Python pipeline in which each module can be executed independently through Jupyter notebooks (exploratory mode) or command-line scripts (pipeline mode). The pipeline does not rely on organism-specific reference models.

### Proteomics data processing and KO mapping

Proteomics data are provided as annotated tables in which each protein entry may be associated with KEGG Orthology (KO) identifiers.^7, 36, 37^ PathwaySeeker parses these inputs and cross-references KO terms with KEGG’s REST API, producing a standardized table (proteomics_with_ko.csv) mapping proteins to KO identifiers.^7, 36, 37^

### Reaction recovery from KO identifiers

For each KO, PathwaySeeker queries the KEGG database to recover all associated reactions.^7, 36, 37^ Redundant or missing entries are handled automatically, and results are compiled into a reaction list (ko_to_reactions.csv).

### Metabolomics annotation and compound mapping

Metabolomics data are provided as feature tables (typically Excel spreadsheets) with metabolites identified or putatively annotated. PathwaySeeker links metabolite names to KEGG compound identifiers (C-numbers).^7, 36, 37^ Because automated annotation is often incomplete, manual curation of metabolite-to-KEGG mappings is encouraged before downstream analysis. The curated table (metabolomics_with_C_numbers_curated.xlsx) ensures higher accuracy in subsequent steps.

### Reaction-to-compound expansion

Reactions derived from proteomics (via KOs) and metabolomics (via C-numbers) are expanded into their constituent substrates and products. PathwaySeeker generates two complementary reaction-compound association files: one from proteomics (reaction_to_compounds_no_cofactors.csv) and one from metabolomics (reaction_to_compounds_from_metabolomics.csv). Common cofactors are optionally removed at this stage.

### Reaction matching and integration

Proteomics- and metabolomics-derived reaction sets are merged into a unified reaction table (matched_metabolites_reactions_all.csv). Reactions are annotated with balanced equations retrieved from KEGG and cached locally to reduce redundant queries.^7, 36, 37^

### Edge generation

Each reaction is converted into directed edges connecting substrates to products via Cartesian product expansion. Multi-substrate or multi-product reactions generate multiple edges. Edge origins are annotated by evidence type.

### Reference Databases

Benchmarking was performed using two KEGG reference sets: KEGG Global – the complete list of known reactions (12,312 reactions), providing a broad measure of recovery across all possible reactions. KEGG *S. cerevisiae* – a species-specific reference (1,512 reactions), providing a more realistic coverage assessment for a model organism.^7, 36, 37^

### Network reconstruction and visualization

The integrated reaction set is represented as a directed bipartite graph using NetworkX. Interactive visualizations are generated with PyVis (graph_all.html), with nodes colored by evidence origin (proteomics, metabolomics, or both) and edges annotated with reaction identifiers and equations. The graph is also exported in JSON format (graph_all.json) for import into interactive exploration module.

### Interactive exploration module

PathwayViz is a Flask wrapper of the open-source Escher software.^38^ In addition to the built-in features of Escher, the Flask application includes backend functions to 1) calculate the positions of the nodes with GraphViz and NetworkX, 2) incorporate additional information such as abundance data from CSV files, and 3) create the JSON files used by Escher to visualize the pathways.^39, 40^ The frontend includes JavaScript and CSS modifications to embed images from PubChem and D3 bar charts, modifying the Escher examples from the Escher Demo GitHub repository (https://github.com/escher/escher-demo).^41, 42^

### Reaction-Level Evaluation

Predicted reactions from PathwaySeeker were compared to the reference sets to compute standard classification metrics: 1. True Positives (TP): predicted reactions present in the reference set; 2. False Positives (FP): predicted reactions that are not present in the reference set; 3. False Negatives (FN): reactions in the reference set not recovered by PathwaySeeker. The metrics were calculated as:

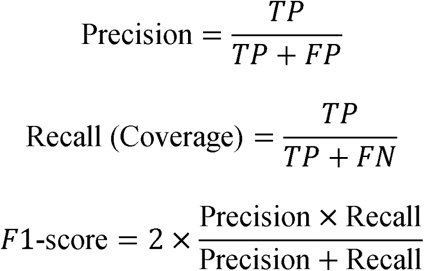

### Integration Metric

Each predicted reaction was assigned an origin based on multi-omics evidence: proteomics only, metabolomics only, or both. Reactions supported by both types of evidence were counted to compute the integration metric:

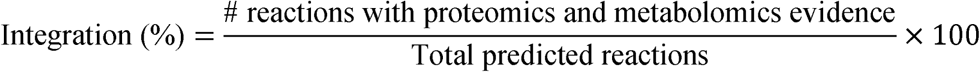

### Equation Correctness

For reactions with KEGG equations available, predicted reaction equations were compared against KEGG entries after normalization (removing prefixes, extra spaces). Exact matches were counted as correct, and the correctness percentage was calculated as:

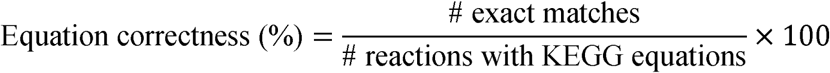

### Compound Coverage

Detected metabolites from experimental metabolomics data were compared to compounds mapped by PathwaySeeker. Coverage was calculated as:

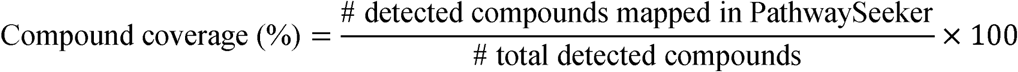

### Cofactor compound set

The following 35 compounds are treated as cofactors and subject to the three-tier constraint strategy described in Methods.

**Water, ions, and gases:** H_2_O, O_2_, CO_2_, NH_3_, H^+^
**Energy and phosphate carriers:** ATP, ADP, AMP, Pi, PPi
**Nucleotides:** UDP, UMP, GMP, GDP, GTP, CTP
**Redox cofactors:** NAD^+^, NADH, NADPH, NADP^+^, FAD, FADH_2_, FMN
**Acyl carriers:** CoA, acetyl-CoA, succinyl-CoA, malonyl-CoA

### Algorithm 1: Oracle-in-the-Loop hypothesis search

The following pseudocode specifies the complete OitL algorithm corresponding to Fig. 1C. Parameters: beam width k = 3, maximum iterations T = 3, convergence threshold theta = 0.70, scoring weights w_e = 0.4, w_c = 0.4, w_p = 0.2.

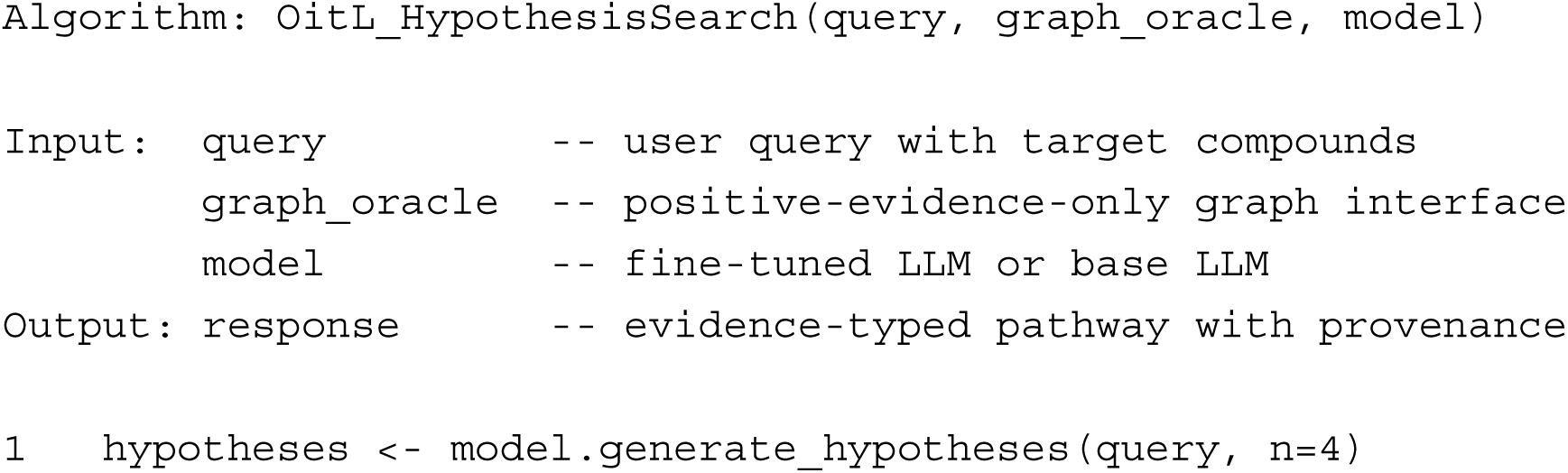
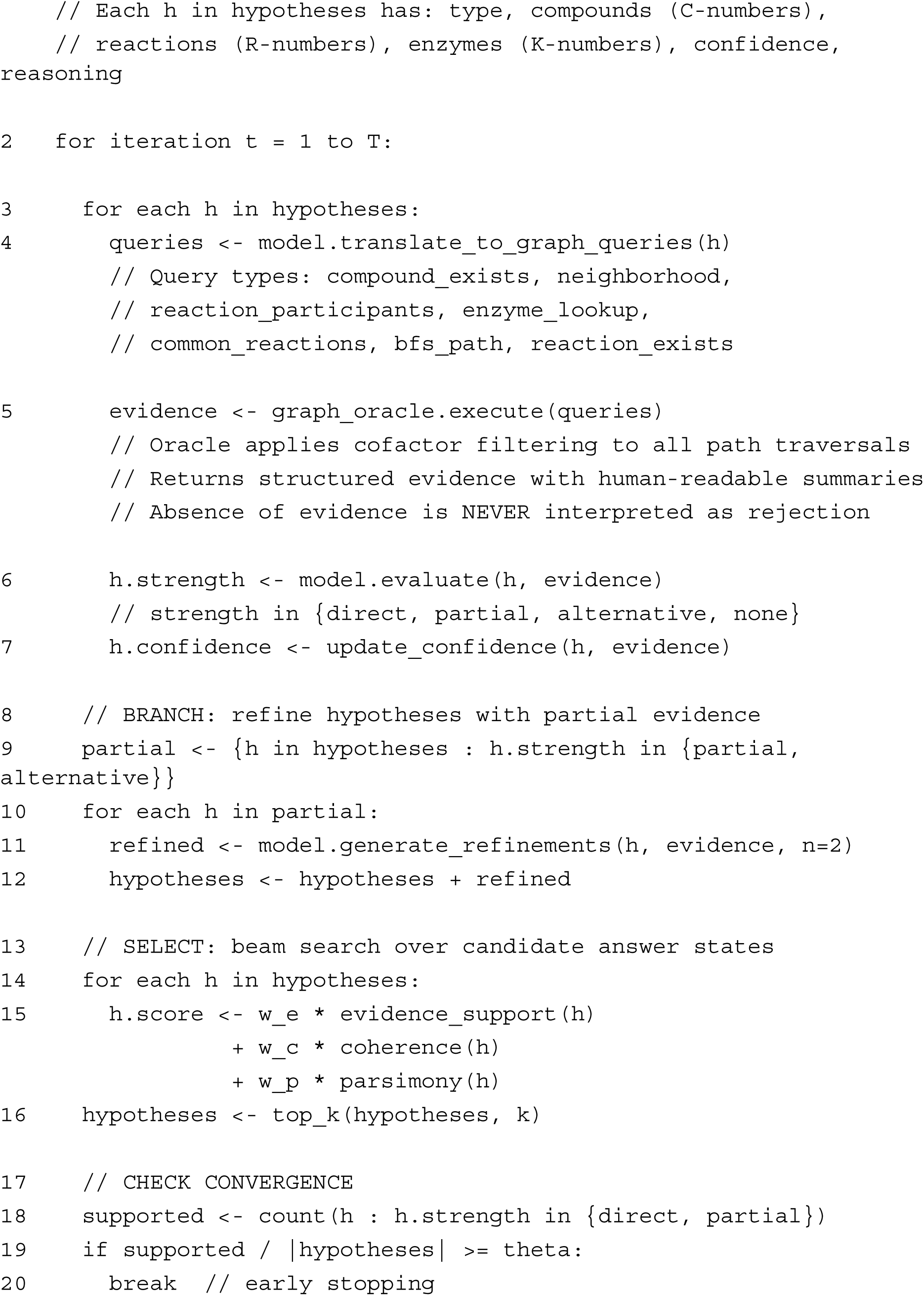
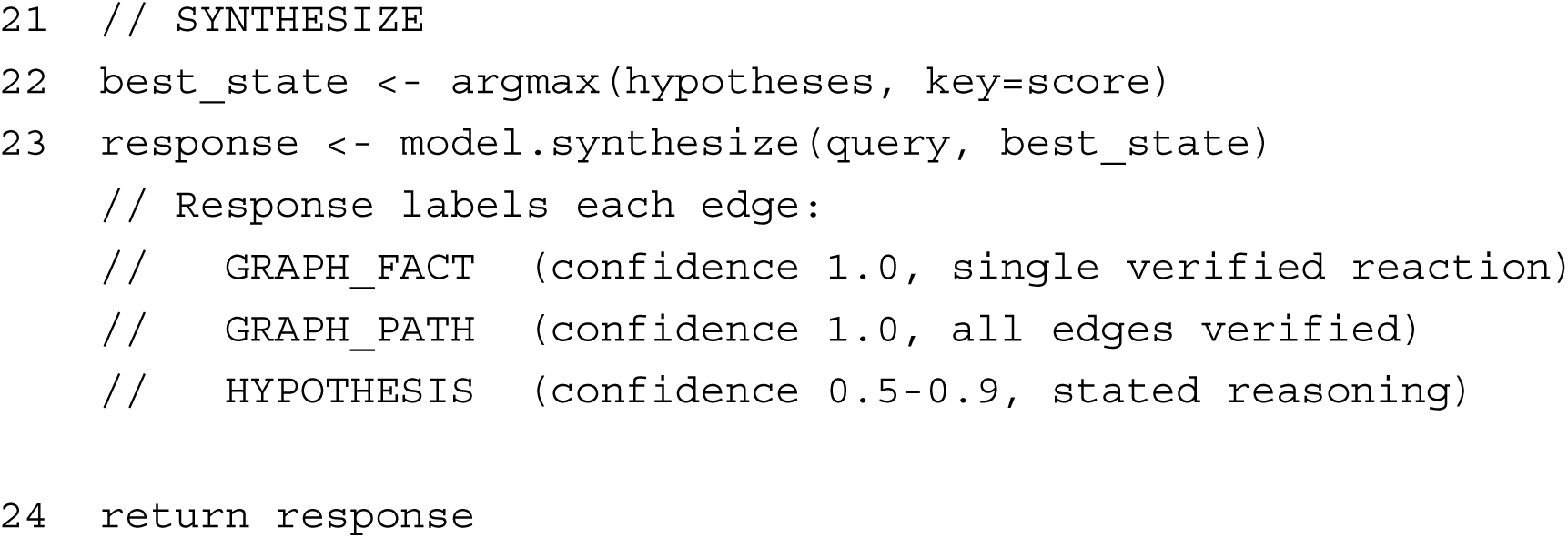

### Graph oracle query types

The graph oracle supports seven query types spanning the compound-reaction-enzyme (C-R-K) schema. All queries enforce the positive-evidence-only principle: results are returned when supporting evidence exists, but absence of a result is never interpreted as biological impossibility.

1. **Compound existence.** Verifies whether a given C-number is present in the reconstructed graph.
2. **Compound neighborhood.** Returns all reactions consuming or producing a given compound, with cofactor nodes excluded from results. Includes the set of non-cofactor neighbors reachable in one reaction step.
3. **Reaction participant lookup.** For a given R-number, returns the substrates, products, and catalyzing enzymes (K-numbers) annotated in the graph.
4. **Enzyme-reaction mapping.** For a given K-number, returns all reactions associated with that enzyme in the graph.
5. **Common reaction identification.** Given a set of compounds, identifies reactions in which two or more of the specified compounds participate as substrates or products.
6. **Breadth-first path search.** Searches for paths between two compounds up to a configurable depth limit (default: 4 reaction steps). Cofactor nodes are excluded from traversal to prevent hub-mediated shortcuts. Returns all shortest paths within the depth limit.
7. **Reaction existence verification.** Verifies whether a given R-number is present in the reconstructed graph.

Each query returns structured evidence comprising a Boolean result, the retrieved data (compounds, reactions, enzymes, or paths as applicable), and a human-readable summary enabling downstream evaluation by the language model.

### Condition-specific metabolic graph construction

PathwaySeeker constructs a knowledge graph from integrated proteomic and metabolomic evidence (**Fig. 1A**). Proteomics data are mapped to KEGG Orthology (KO) identifiers and expanded to associated reactions via the KEGG REST API. Metabolomics data are mapped to KEGG compound identifiers (C-numbers) through automated annotation followed by manual curation. The two reaction sets are merged using an inclusive, OR-based integration logic: a reaction is retained if supported by either proteomic evidence (a detected enzyme catalyzing that reaction) or metabolomic evidence (detected substrates or products), without requiring simultaneous detection across both omics layers. Each edge in the resulting graph is annotated with its evidence source (proteomics, metabolomics, or both). Reconstructed reactions were benchmarked against two KEGG reference sets: the global KEGG reaction universe (12,312 reactions) and the *Saccharomyces cerevisiae* reaction set (1,512 reactions).

### Schema-aware training data generation

Training data were generated programmatically from the reconstructed graph using a schema-aware sampling approach adapted from the Graph-aligned Language Model (GLaM) framework (**Fig. 1B**).^27^ Rather than sampling arbitrary graph paths, we defined the fundamental unit of metabolic meaning as a transformation unit: a set of substrates converted to products via a specific reaction catalyzed by one or more enzymes, respecting the three-layer compound-reaction-enzyme (C-R-K) structure of metabolic networks.

The generator produces examples across five evidence types, each teaching the model a distinct reasoning behavior. GRAPH_FACT examples assert single verified reactions (e.g., ‘What are the substrates and products of R00697?’, the phenylalanine ammonia-lyase reaction), grounding the model in reaction-level knowledge at confidence 1.0. GRAPH_PATH examples describe multi-step chains in which all edges are experimentally confirmed (e.g., the four-reaction sequence from L-phenylalanine through trans-cinnamate, 4-coumarate, and caffeate to ferulate), teaching the model to trace verified pathways while distinguishing them from merely reachable connections. HYPOTHESIS examples describe connections that are biochemically plausible but absent from the graph (e.g., a candidate route bridging two disconnected components via an undetected intermediate), with confidence scaled according to the strength of supporting neighborhood evidence and stated biochemical reasoning. NO_PATH examples present compound pairs located in disconnected graph components, teaching the model to recognize the absence of verified connectivity while maintaining that such absence reflects incomplete experimental coverage rather than biological impossibility. INVALID examples present queries that violate metabolic constraints (e.g., ‘Is there a pathway from ATP to ferulate?’), training the model to reject cofactor-endpoint requests and cofactor-mediated shortcuts even when they are technically reachable in the graph.

Each training example follows a structured chat-completion format: a system prompt establishing the model as a graph-constrained metabolic reasoning assistant, a user prompt containing serialized local graph context and the query, and an assistant response in JSON specifying the evidence type, answer, and confidence score. This format enables automated verification of model outputs against the ground-truth graph.

### Cofactor constraint strategy

Cofactors and ubiquitous metabolites present a fundamental challenge for graph-based pathway inference. Compounds such as ATP, NAD(P)H, and CoA participate in hundreds of reactions, creating artificial shortcuts that connect metabolically unrelated compounds through highly connected hubs. To address this, we implemented a three-tier constraint strategy:

1. **Endpoint exclusion**: Cofactors are disallowed as start or end nodes in predicted pathways. Biologically meaningful pathway queries should define objectives in terms of core metabolites, not energy or redox currencies.
2. **Inferred edge restriction**: Cofactors may participate only in verified reactions supported by existing annotations. When proposing hypothetical connections (HYPOTHESIS evidence type), inferred edges must not be cofactor-mediated, preventing the model from using hub nodes to bridge disconnected pathway components.
3. **Penalty-based weig**hting: During path search, edges involving cofactors incur a high traversal penalty (default: 10.0x), ensuring they are used only when mechanistically unavoidable. This preserves biochemical realism while enabling the model to recognize verified cofactor participation in specific reactions.

The cofactor set comprises 35 compounds including water, ions, and gases (H_2_O, O_2_, CO_2_, NH_3_, H^+^); energy and phosphate carriers (ATP, ADP, AMP, pi, PPi); nucleotides (UDP, UMP, GMP, GDP, GTP, CTP); redox cofactors (NAD^+^, NADH, NADPH, NADP^+^, FAD, FADH_2_, FMN); and acyl carriers (CoA, acetyl-CoA, succinyl-CoA, malonyl-CoA).

### Training data balance and model fine-tuning

The reconstructed *T. versicolor* network contains approximately 1.3 million possible non-cofactor compound pairs (1,153 backbone metabolites), of which only 2,831 (0.21%) are connected by verified metabolic paths, across 1,828 disconnected components. To prevent imbalanced training (>60% negative examples under naive sampling), we exhaustively extracted all verifiable positive paths before negative sampling and imposed hard caps on negative examples (20% or less of total). The final training set of 16,422 examples achieved 74% positive examples (GRAPH_FACT, GRAPH_PATH, and HYPOTHESIS) with 20% negative (NO_PATH and INVALID). PathwaySeeker was fine-tuned from GPT-4.1 on Azure OpenAI with batch size 8, learning rate multiplier 1.2, and 3 epochs, yielding approximately 6,150 gradient steps across 49,000 example passes.

### Oracle-in-the-Loop hypothesis search

The central inference challenge is that the experimental graph is incomplete: the *T. versicolor* network comprises 19 disconnected components, and any single query may span verified edges, partially supported connections, and regions where the graph coverage is missing. A single-pass generation from the fine-tuned model cannot reliably distinguish these cases, because the model’s biochemical priors will fill gaps without signaling that it has left the experimental evidence. The Oracle-in-the-Loop algorithm addresses this by interleaving generation with verification (**Fig. 2**).

At each step, the model proposes candidate connections, the graph oracle checks them against experimental evidence, and the model revises its hypotheses based on what the oracle returns. This iterative structure ensures that the final output reflects the graph’s actual evidence landscape rather than the model’s prior expectations. Two feedback loops govern the search. The inner loop (Branch) refines individual hypotheses that receive partial graph support, directing the model to explore evidence-suggested routes rather than relying solely on biochemical reasoning. The outer loop (Select) operates at the level of the full candidate set, applying beam search with composite scoring to balance evidence support, biochemical coherence, and parsimony across competing pathway explanations. Early stopping prevents unnecessary iteration when the evidence is decisive. The algorithm is architecturally modular: it accepts either the fine-tuned PathwaySeeker model or a base LLM as the underlying reasoning engine.

### Graph oracle

The graph oracle enforces the positive-evidence-only principle: queries return supporting evidence when present, but absence is never interpreted as rejection. It supports seven query types (compound existence, neighborhood retrieval with cofactor exclusion, reaction participant lookup, enzyme-reaction mapping, common reaction identification, breadth-first path search with configurable depth limit, and reaction existence verification), each returning structured evidence with human-readable summaries for downstream evaluation by the language model.

### Evaluation framework

#### Experimental Evidence Ratio (EER)

EER quantifies the provenance composition of each response as the fraction of reported pathway edges directly confirmed by the organism-specific graph: EER = (edges with GRAPH_FACT or GRAPH_PATH evidence) / (total edges in response).

#### LLM-as-judge quality assessment

An independent language model (GPT-4.1) scored each response on four dimensions (1 to 5 scale): Scientific Reasoning, Specificity, Evidence Transparency, and Clarity. The evaluator received only the query and response, without access to evaluation mode or system configuration.

#### Expert blind evaluation

Domain scientists evaluated paired responses (graph-verified-only versus hypothesis mode) in blinded format, selecting their preferred response with written justification.

### Thermodynamic feasibility analysis

Max-min Driving Force (MDF) analysis ^34^ was performed using equilibrator_api (v0.4.5) and equilibrator_pathway (v0.4.4) at pH 7.0, ionic strength 250 mM, pMg 3.0, with metabolite concentrations constrained between 0.00001 and 10 mM.

## Acknowledgements.

This research was supported by the Environmental Molecular Sciences Laboratory, a DOE Office of Science User Facility sponsored by the Biological and Environmental Research program under Contract No. DE-AC05-76RL01830.

## Author Contributions

Project Conceptualization: LMOM, SC; Experimental Investigation: LMOM, NBC, MTO, SC; Analytical Data Collection: LMOM; Data Curation, Interpretation, Analysis: LMOM, NBC, MTO, SC; Software–Original Code: LMOM, NBC, MTO, SC; Writing– Original Draft: LMOM, SC; Writing– Review & Editing: LMOM, NBC, MTO, SC, JEM, KGS, JPB; Supervision: JEM, KGS, JPB.

All authors read and approved the manuscript.

## Data availability statement

The complete codebase, including graph construction, training data generation, OitL search, and evaluation modules: https://github.com/pnnl/PathwaySeeker. All data utilized in the manuscript and generated throughout the project is provided in the repository, manuscript and supplemental files.

## References

1. Bordbar, A., Monk, J.M., King, Z.A. & Palsson, B.O. Constraint-based models predict metabolic and associated cellular functions. Nature Reviews Genetics 15, 107–120 (2014).

2. Chubukov, V., Gerosa, L., Kochanowski, K. & Sauer, U. Coordination of microbial metabolism. Nature Reviews Microbiology 12, 327–340 (2014).

3. Nielsen, J. & Keasling, J.D. Engineering cellular metabolism. Cell 164, 1185–1197 (2016).

4. Azam, M. et al. A comprehensive evaluation of large language models in mining gene relations and pathway knowledge. Quantitative Biology 12, 360–374 (2024).

5. M. Bran, A., et al. Augmenting large language models with chemistry tools. Nature machine intelligence 6, 525–535 (2024).

6. Roohani, Y. et al. Biodiscoveryagent: An ai agent for designing genetic perturbation experiments. arXiv preprint arXiv:2405.17631 (2024).

7. Kanehisa, M. & Goto, S. KEGG: Kyoto Encyclopedia of Genes and Genomes. Nucleic Acids Research 28, 27–30 (2000).

8. Karp, P.D. et al. The BioCyc collection of microbial genomes and metabolic pathways. Briefings in bioinformatics 20, 1085–1093 (2019).

9. Caspi, R. et al. The MetaCyc database of metabolic pathways and enzymes-a 2019 update. Nucleic acids research 48, D445–D453 (2020).

10. Wishart, D.S. et al. HMDB 5.0: the human metabolome database for 2022. Nucleic acids research 50, D622–D631 (2022).

11. Thiele, I. & Palsson, B.Ø. A protocol for generating a high-quality genome-scale metabolic reconstruction. Nature protocols 5, 93–121 (2010).

12. Fang, X., Lloyd, C.J. & Palsson, B.O. Reconstructing organisms in silico: genome-scale models and their emerging applications. Nature Reviews Microbiology 18, 731–743 (2020).

13. Monteiro, L.M.O. et al. Metabolic profiling of two white-rot fungi during 4-hydroxybenzoate conversion reveals biotechnologically relevant biosynthetic pathways. Communications Biology 8, 224 (2025).

14. Subramanian, I., Verma, S., Kumar, S., Jere, A. & Anamika, K. Multi-omics data integration, interpretation, and its application. Bioinformatics and biology insights 14, 1177932219899051 (2020).

15. Reel, P.S., Reel, S., Pearson, E., Trucco, E. & Jefferson, E. Using machine learning approaches for multi-omics data analysis: A review. Biotechnology advances 49, 107739 (2021).

16. Misra, B.B., Langefeld, C., Olivier, M. & Cox, L.A. Integrated omics: tools, advances and future approaches. Journal of molecular endocrinology 62, R21–R45 (2019).

17. Di Filippo, M. et al. INTEGRATE: Model-based multi-omics data integration to characterize multi-level metabolic regulation. PLoS computational biology 18, e1009337 (2022).

18. Töpfer, N., Kleessen, S. & Nikoloski, Z. Integration of metabolomics data into metabolic networks. Frontiers in plant science 6, 49 (2015).

19. Hackett, S.R. et al. Systems-level analysis of mechanisms regulating yeast metabolic flux. Science 354, aaf2786 (2016).

20. Zhang, H., et al. mosGraphGPT: a foundation model for multi-omic signaling graphs using generative AI. bioRxiv (2024).

21. Guo, Z., Duan, D., Liang, Y., Patil, A. & Xie, P. MetaboliteChat: A Unified Multimodal Large Language Model for Interactive Metabolite Analysis and Functional Insights. bioRxiv, 2025.2011. 2007.687008 (2025).

22. Arowolo, M.O. et al. Agentic RAG-Driven Multi-Omics Analysis for PI3K/AKT Pathway Deregulation in Precision Medicine. Algorithms 18, 545 (2025).

23. Wei, J. et al. Chain-of-thought prompting elicits reasoning in large language models. Advances in neural information processing systems 35, 24824–24837 (2022).

24. Sprueill, H., et al. in Findings of the Association for Computational Linguistics: EMNLP 2023 8348-8365 (2023).

25. Sprueill, H.W., et al. ChemReasoner: Heuristic search over a large language model’s knowledge space using quantum-chemical feedback. arXiv preprint arXiv:2402.10980 (2024).

26. Hao, S. et al. in Proceedings of the 2023 Conference on Empirical Methods in Natural Language Processing 8154-8173 (2023).

27. Dernbach, S., Agarwal, K., Zuniga, A., Henry, M., & Choudhury, S. GLaM: Fine-Tuning Large Language Models for Domain Knowledge Graph Alignment via Neighborhood Partitioning and Generative Subgraph Encoding. AAAI Spring Symposium: AAAI-MAKE 2024 (2024).

28. Lightman, H., et al. in The twelfth international conference on learning representations (2023).

29. Yao, S. et al. Tree of thoughts: Deliberate problem solving with large language models. Advances in neural information processing systems 36, 11809–11822 (2023).

30. Kijpornyongpan, T., Schwartz, A., Yaguchi, A. & Salvachúa, D. Systems biology-guided understanding of white-rot fungi for biotechnological applications: A review. iScience 25, 104640 (2022).

31. del Cerro, C. et al. Intracellular pathways for lignin catabolism in white-rot fungi. Proceedings of the National Academy of Sciences 118, e2017381118 (2021).

32. Kuatsjah, E., et al. Biochemical and structural characterization of enzymes in the 4-hydroxybenzoate catabolic pathway of lignin-degrading white-rot fungi. Cell Reports 43 (2024).

33. Reed, J.L. et al. Systems approach to refining genome annotation. Proceedings of the National Academy of Sciences 103, 17480–17484 (2006).

34. Beber, M.E. et al. eQuilibrator 3.0: a database solution for thermodynamic constant estimation. Nucleic Acids Research 50, D603–D609 (2021).

35. Kim, J. et al. Multi-Omics Driven Metabolic Network Reconstruction and Analysis of Lignocellulosic Carbon Utilization in Rhodosporidium toruloides. Frontiers in Bioengineering and Biotechnology Volume 8 - 2020 (2021).

36. Kanehisa, M. Toward understanding the origin and evolution of cellular organisms. Protein science 28, 1947–1951 (2019).

37. Kanehisa, M., Furumichi, M., Sato, Y., Matsuura, Y. & Ishiguro-Watanabe, M. KEGG: biological systems database as a model of the real world. Nucleic acids research 53, D672–D677 (2025).

38. King, Z.A. et al. Escher: a web application for building, sharing, and embedding data-rich visualizations of biological pathways. PLoS computational biology 11, e1004321 (2015).

39. Ellson, J., Gansner, E., Koutsofios, L., North, S.C. & Woodhull, G. in International symposium on graph drawing 483–484 (Springer, 2001).

40. Hagberg, A., Swart, P.J. & Schult, D.A. (Los Alamos National Laboratory (LANL), 2007).

41. Bostock, M., Ogievetsky, V. & Heer, J. D³ data-driven documents. IEEE transactions on visualization and computer graphics 17, 2301–2309 (2011).

42. Kim, S., et al. PubChem 2025 update. Nucleic Acids Research 53, D1516–D1525 (2024).

