## supplementary file for "PathwaySeeker: Evidence-Grounded AI Reasoning over Organism-Specific Metabolic Networks"

^1^ Biological Sciences Division, Pacific Northwest National Laboratory, Richland Washington
^2^ AI & Data Analytics, Pacific Northwest National Laboratory, Richland Washington  ^3^ Environmental Molecular Science Division, Pacific Northwest National Laboratory, Richland Washington ^4^ Department of Molecular Microbiology and Immunology, Oregon Health & Science University, Portland Oregon  ^5^ **Advanced Comp, Math & Data Division,** Pacific Northwest National Laboratory, Richland Washington

### Supplementary text

#### Agentic architecture and distribution model

PathwaySeeker is implemented as three modular agents: a Graph Builder that reconstructs the organism-specific metabolic network from integrated multi-omics evidence, an Oracle-in-the-Loop Hypothesis Search agent that executes evidence-typed pathway inference via the seven-stage beam search algorithm, and a Query Extractor that translates natural-language queries into structured graph operations. These agents are designed for distribution as Model Context Protocol (MCP) server connectors, enabling researchers to attach organism-specific metabolic reasoning to general-purpose AI assistants without installing or configuring a standalone computational pipeline. This distribution model lowers the adoption barrier from computational proficiency to domain expertise: a researcher with proteomic and metabolomic data from any organism with sufficient coverage can reconstruct an organism-specific graph and begin querying it conversationally, without requiring curated genome-scale models or prior pathway annotations. The complete codebase, including graph construction, training data generation, OitL search, and evaluation modules, is available at [GitHub URL].

#### Factors governing system performance

**Base model capability and the persistence of the grounding gap**. Comparison of PathwaySeeker's outputs against those of ungrounded general-purpose models reveals that increasing base model capability does not resolve the fundamental grounding problem. GPT-4.1 without graph access produces biochemically accurate but ungrounded pathway descriptions; it cannot identify which reactions are supported by the user's experimental data. GPT-5.2, whose training corpus includes Monteiro et al. (2025),^1^ returns the correct pathway architecture by citing the published literature, yet still cannot distinguish claims derived from the user's experiment from those absorbed during pretraining. As frontier models improve and incorporate more published results, they will generate increasingly convincing responses that are increasingly difficult to audit for experimental grounding, making explicit evidence provenance more necessary, not less. The grounding gap is structural: it arises from the absence of runtime verification against condition-specific evidence, not from insufficient model capability.

**Contribution of fine-tuning versus search architecture**. PathwaySeeker's OitL algorithm is architecturally modular: it accepts either the fine-tuned model or a base language model as the reasoning engine. The search architecture alone, comprising the graph oracle, positive-evidence-only retrieval, evidence-typed output format, and hypothesis-driven beam search, enforces experimental grounding at every reasoning step regardless of the underlying model's training. Fine-tuning adds a complementary benefit: the model internalizes the graph's topology and evidence structure, producing higher evidence density and greater specificity in its initial hypotheses, which in turn reduces the number of search iterations required for convergence. For prospective adopters, this decomposition implies that the Oracle-in-the-Loop pattern provides immediate value with any capable base model, while organism-specific fine-tuning represents an incremental investment with measurable returns in output quality.

#### Benchmarking PathwaySeeker Predictions

Quantitatively, PathwaySeeker generated a comprehensive mapping of metabolic reactions, compounds, and edges for T. versicolor. Across the dataset, the tool predicted **3,402 unique reactions**, connecting **1,192 compounds** through **1,897 directed edges** (substrate -> product). Importantly, **all predicted reaction equations perfectly matched KEGG references (**100% equation correctness), demonstrating the tool’s reproducibility and reliability in reaction balancing. When compared to the global KEGG universe (12,312 reactions), PathwaySeeker achieved **perfect precision (1.0),** indicating no false-positive reactions, with a **recall of 27.6%,** reflecting the expected partial coverage for a non-model organism. The **F1-score was 0.433,** highlighting a balanced combination of coverage and precision. Relative to Saccharomyces cerevisiae (1,512 KEGG reactions), recall improved to **72.6%,** confirming the ability of PathwaySeeker to recover organism-specific pathways.

When benchmarked against the global KEGG universe, PathwaySeeker achieved **perfect precision** with moderate coverage (27.6%), reflecting expected partial recovery in a non-model organism. Comparison to S. cerevisiae reactions showed **higher recall (72.6%)** but lower precision (0.323), as some predicted reactions are not present in the yeast-specific reference. This dual benchmarking demonstrates PathwaySeeker’s ability to recover biologically relevant reactions while remaining consistent with organism-specific metabolism.

Integration of multi-omics evidence was also quantified: **26% of edges (493/1,897) were supported by both proteomics and metabolomics data**, while **54.4% were proteomics-only** and **19.6% metabolomics-only,** providing insight into confidence and data convergence. Compound mapping further confirmed high coverage: of 181 metabolites detected in metabolomics experiments, **163 were correctly mapped by PathwaySeeker,** corresponding to **90.1% compound coverage.**

Overall, PathwaySeeker delivers **accurate, reproducible, and quantitatively benchmarked networks** with automated reaction-to-compound mapping, multi-omics integration, and fully balanced reactions, all without requiring a species-specific model. The tool completed network reconstruction efficiently, leveraging cached KEGG equations to minimize redundant queries, thereby providing a robust framework for hypothesis generation in non-model fungal systems.

The tool successfully reconstructed a coherent sequence of reactions corresponding to the canonical early phenylpropanoid pathway: L-phenylalanine (C00079) -> trans-cinnamate (C00423) -> p-coumarate (C00811) -> caffeate (C01197) -> ferulate (C01494). Each transition was automatically mapped to validated enzymatic reactions - phenylalanine ammonia-lyase (PAL, EC 4.3.1.24), cinnamate 4-hydroxylase (C4H, EC 1.14.14.91), p-coumarate 3-hydroxylase, and caffeate O-methyltransferase (COMT, EC 2.1.1.68) - without prior model constraints.

Importantly, PathwaySeeker identified this pathway **without prior knowledge or species-specific models,** automating what previously required manual curation. The resulting network not only captures canonical reactions but also integrates proteomics-derived evidence, providing a reproducible and interactive framework for exploring lignin-related metabolism in non-model fungi. These results illustrate PathwaySeeker’s ability to recover biologically plausible, multi-step pathways and highlight its potential to accelerate hypothesis generation in organisms lacking curated genome-scale models. These results illustrate that PathwaySeeker supports hypothesis generation in systems lacking curated genome-scale models, offering transparent integration of multi-omics evidence into metabolic networks.

Overall, PathwaySeeker delivers **accurate, reproducible, and quantitatively benchmarked networks** with automated reaction-to-compound mapping, multi-omics integration, and fully balanced reactions, all without requiring a species-specific model. Cached KEGG equations minimized redundant queries, enabling efficient network reconstruction and supporting rapid hypothesis generation in non-model fungal systems.

### Supplementary Figures

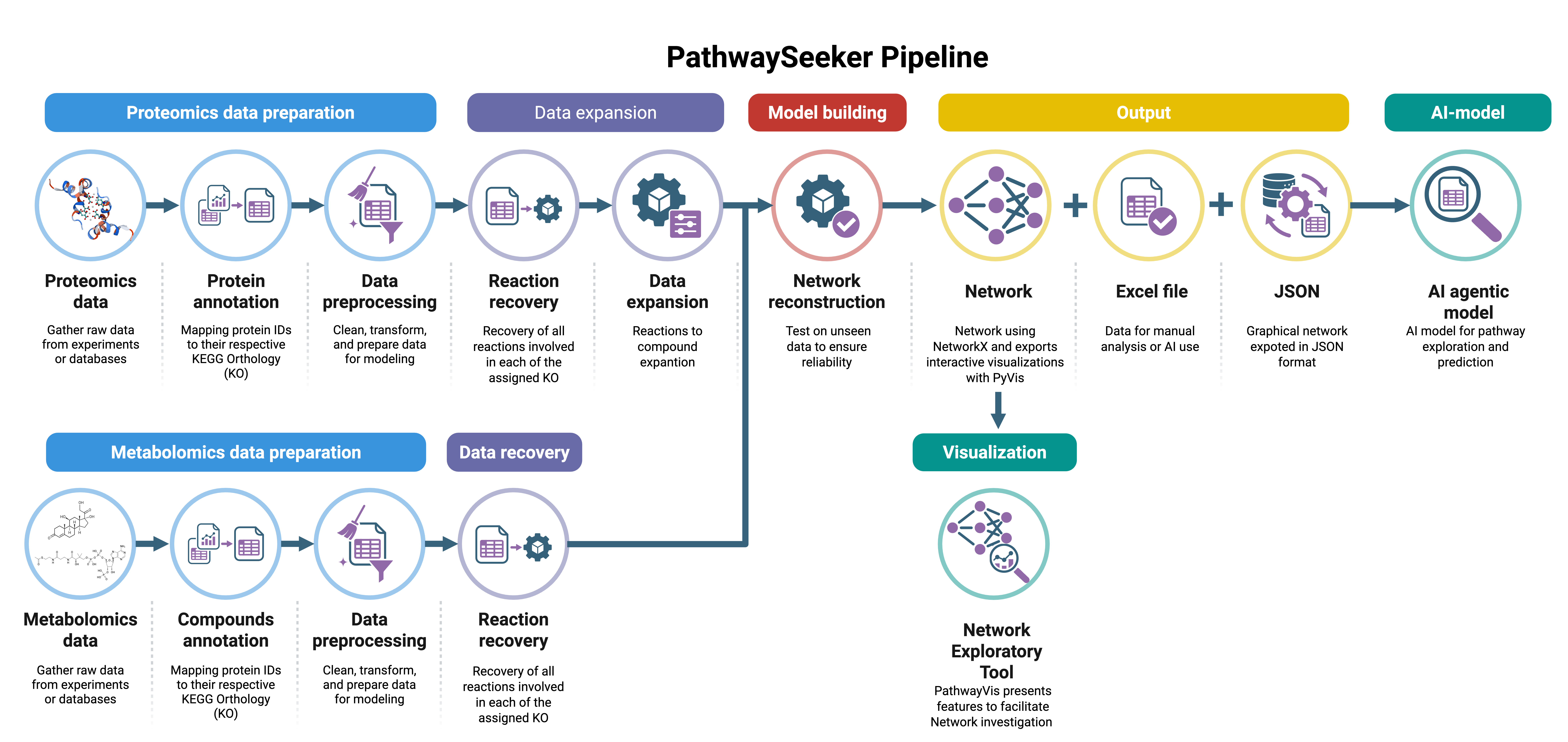

Supplementary fig 1. PathwaySeeker AI-guided framework for integrative multi-omics pathway reconstruction and hypothesis generation.

PathwaySeeker implements a modular computational architecture that integrates proteomic and metabolomic datasets into a unified, reaction-level knowledge graph. The workflow comprises (i) data preparation and harmonization, (ii) graph-grounded metabolic network reconstruction, and (iii) AI-assisted pathway expansion and hypothesis generation. Experimental evidence is encoded at the reaction level to distinguish graph-supported routes from inferred extensions. The system produces interpretable outputs, including reconstructed pathway files, annotated reaction networks, and interactive visualizations. An integrated AI module operates on the structured graph representation to propose evidence-typed pathway hypotheses grounded in biochemical topology.

### Supplementary tables

Several tools have been developed for multi-omics data analysis, yet they present important limitations. **MetaboAnalyst**, for instance, supports metabolomics, transcriptomics, and gene/KO inputs and includes a Network Explorer module for visual exploration or Debiased Sparse Partial Correlation (DSPC) network analysis. Nevertheless, these networks remain correlation-based, lacking balanced reaction reconstruction or mechanistic integration of proteomics and metabolomics. **STRING** and the **Cytoscape StringApp** focus on protein–protein interaction networks, but they are largely restricted to model organisms and do not reconstruct reactions. Similarly, **OmicsNet**, **PASNet**, and **iOmicsPASS** provide multi-omics integration but rely on associations or machine learning rather than explicit biochemical reactions, often requiring predefined species models. Tools such as **MetScape** and **ReactomeGSA** enable partial integration of metabolomics and transcriptomics but are still oriented toward enrichment or visualization rather than pathway reconstruction. Finally, frameworks like **MOFA+** and **MixOmics** are powerful for factor analysis and cross-omics correlations, yet they are not designed for reconstructing biochemical routes.

#### Supplementary Table 1. List of commonly used multi-omic tools

| **Tool** | **Supported data types** | **Non-model organisms** | **Pathway reconstruction** | **Multi-omics integration** | **Visualization / output** | **Ref.** |
| --- | --- | --- | --- | --- | --- | --- |
| **PathwaySeeker** | Proteomics, Metabolomics | Yes | Yes (KEGG-based, balanced reactions) | Yes (direct mapping + network) | Tables, JSON, interactive HTML graphs | This manuscript |
| **MetaboAnalyst** | Metabolomics | Partial ^a^ | No (does not reconstruct balanced reactions) | Yes (limited, correlation-based) | Web interface, charts, pathway maps | ^2^ |
| **OmicsNet** | Proteomics, Metabolomics, Transcriptomics | Partial ^a^ | No | Yes | Network visualization (Cytoscape-based) | ^3^ |
| **STRING** | Proteomics, Transcriptomics | Partial ^a^ | No (associations only) | No | Network visualization, web interface | ^4^ |
| **PASNet** | Proteomics, Transcriptomics | Partial ^a^ | No | Yes (deep learning) | Graphs / network models | ^5^ |
| **MS2MP** | Metabolomics | Partial ^a^ | No | No | Web interface, pathway visualization | ^6^ |
| **Cytoscape StringApp** | Proteomics, Transcriptomics | Partial ^a^ | No (associations only) | No | Cytoscape network visualization | ^7^ |
| **OmicsIntegrator** | Proteomics, Transcriptomics | Yes | Partial (context-specific networks) | Yes | Network output, Cytoscape compatible | ^8^ |
| **iOmicsPASS / iOmicsPath** | Proteomics, Metabolomics, Transcriptomics | Partial ^a^ | Partial (pathway scoring) | Yes | Network visualization | ^9^ |
| **MetScape (Cytoscape app)** | Metabolomics, Transcriptomics | Partial ^a^ | Partial (metabolic network mapping) | Yes | Cytoscape visual networks | ^10, 11^ |
| **ReactomeGSA** | Transcriptomics, Proteomics | Partial ^a^ | Partial (pathway enrichment) | Partial | Web-based pathway-level visualization | ^12^ |
| **MOFA+** | Transcriptomics, Proteomics, Metabolomics | Yes | No | Yes (latent factors across omics) | R/Python plots, factor heatmaps | ^13^ |
| **MixOmics / MONI** | Various (multi-omics) | Partial ^a^ | No | Yes (network inference or dimension reduction) | R plots, network representation | ^14^ |

^a^ The application presents a few non-model species genomes available

#### Supplementary ****Table 2. Edge origins and multi-omics integration****

| **Edge origin** | **Number of edges** | **% of total** |
| --- | --- | --- |
| Proteomics only | 1,032 | 54.4% |
| Metabolomics only | 372 | 19.6% |
| Both (integration) | 493 | 26.0% |
| Unknown | 0 | 0.0% |

Note: Each edge represents a substrate → product relation derived from a reaction.

#### Supplementary ****Table 3. Compound coverage****

| **Metric** | **Value** |
| --- | --- |
| Metabolomics-detected compounds | 181 |
| Compounds mapped by PathwaySeeker | 1,192 |
| Overlap (detected ∩ mapped) | 163 |
| Compound coverage (%) | 90.1% |

#### Supplementary Table 4. Reaction-level summary (KEGG global vs. S. cerevisiae)

| **Metric** | **KEGG Global** | **KEGG S. cerevisiae** |
| --- | --- | --- |
| Unique predicted reactions | 3,402 | 3,402 |
| KEGG reactions | 12,312 | 1,512 |
| True positives (TP) | 3,402 | 1,097 |
| False positives (FP) | 0 | 2,305 |
| False negatives (FN) | 8,910 | 415 |
| Precision | 1.0 | 0.323 |
| Recall (coverage) | 27.6% | 72.6% |
| F1-score | 0.433 | 0.447 |
| Equation correctness (%) | 100% | 100% |

Note: Coverage and precision are computed relative to the respective KEGG reference.

### References

1.	Monteiro, L.M.O. et al. Metabolic profiling of two white-rot fungi during 4-hydroxybenzoate conversion reveals biotechnologically relevant biosynthetic pathways. *Communications Biology* **8**, 224 (2025).

2.	Pang, Z. et al. Using MetaboAnalyst 5.0 for LC–HRMS spectra processing, multi-omics integration and covariate adjustment of global metabolomics data. *Nature Protocols* **17**, 1735-1761 (2022).

3.	Zhou, G. & Xia, J. OmicsNet: a web-based tool for creation and visual analysis of biological networks in 3D space. *Nucleic Acids Research* **46**, W514-W522 (2018).

4.	Szklarczyk, D. et al. The STRING database in 2023: protein–protein association networks and functional enrichment analyses for any sequenced genome of interest. *Nucleic Acids Research* **51**, D638-D646 (2022).

5.	Hao, J., Kim, Y., Kim, T.-K. & Kang, M. PASNet: pathway-associated sparse deep neural network for prognosis prediction from high-throughput data. *BMC Bioinformatics* **19**, 510 (2018).

6.	Bao, H. et al. MS2MP: A Deep Learning Framework for Metabolic Pathway Prediction from MS/MS-Based Untargeted Metabolomics. *Analytical Chemistry* **97**, 14200-14209 (2025).

7.	Doncheva, N.T., Morris, J.H., Gorodkin, J. & Jensen, L.J. Cytoscape StringApp: Network Analysis and Visualization of Proteomics Data. *Journal of Proteome Research* **18**, 623-632 (2019).

8.	Tuncbag, N. et al. Network-Based Interpretation of Diverse High-Throughput Datasets through the Omics Integrator Software Package. *PLOS Computational Biology* **12**, e1004879 (2016).

9.	Koh, H.W.L. et al. iOmicsPASS: network-based integration of multiomics data for predictive subnetwork discovery. *npj Systems Biology and Applications* **5**, 22 (2019).

10.	Basu, S. et al. Sparse network modeling and metscape-based visualization methods for the analysis of large-scale metabolomics data. *Bioinformatics* **33**, 1545-1553 (2017).

11.	Karnovsky, A. et al. Metscape 2 bioinformatics tool for the analysis and visualization of metabolomics and gene expression data. *Bioinformatics* **28**, 373-380 (2011).

12.	Grentner, A. et al. ReactomeGSA: new features to simplify public data reuse. *Bioinformatics* **40** (2024).

13.	Argelaguet, R. et al. MOFA+: a statistical framework for comprehensive integration of multi-modal single-cell data. *Genome Biology* **21**, 111 (2020).

14.	Rohart, F., Gautier, B., Singh, A. & Lê Cao, K.-A. mixOmics: An R package for ‘omics feature selection and multiple data integration. *PLOS Computational Biology* **13**, e1005752 (2017).
